# Constructing threat probability, fear behaviour, and aversive prediction error in the brainstem

**DOI:** 10.1101/2021.11.19.469307

**Authors:** Jasmin A. Strickland, Michael A. McDannald

## Abstract

When faced with potential threat we must estimate its probability, respond advantageously, and leverage experience to update future estimates. Threat estimates are the proposed domain of the forebrain, while behaviour is elicited by the brainstem. Yet, the brainstem is also a source of prediction error, a learning signal to acquire and update threat estimates. Neuropixels probes allowed us to record single-unit activity across a 21-region brainstem axis during probabilistic fear discrimination with foot shock outcome. Against a backdrop of diffuse behaviour signaling, a brainstem network with a dorsal hub signaled threat probability. Neuronal function remapping during the outcome period gave rise to brainstem networks signaling prediction error and shock on multiple timescales. The results reveal construction of threat probability, behaviour, and prediction error along a single brainstem axis.

**One-Sentence Summary:** The brainstem constructs threat probability, behaviour, and prediction error from neuronal building blocks.

## Introduction

Faced with potential threat, we must estimate its probability, determine an appropriate response, and – should we come away intact – adjust our estimates for future encounters. Historical and current descriptions of the brain’s threat circuitry emphasize a division of labour in which forebrain regions estimate threat probability, while the brainstem elicits behaviour (*1, 2*). However, behaviour signaling is not observed in expected brainstem neuronal populations, such as the periaqueductal gray (*3, 4*), which instead signals threat probability (*5*). Instead, periaqueductal behaviour signaling is observed in an unexpected population (*6*). Further, the brainstem periaqueductal gray is a source of prediction error (*3, 7, 8*), a learning signal to adjust threat estimates (*9*). These findings necessitate a more complex role for the brainstem in threat estimation. However, evidence of widespread brainstem threat probability signaling remains elusive, and more complete descriptions of brainstem behaviour and prediction error signaling are needed. Recording a 21-region axis with Neuropixels (*10*) during probabilistic fear discrimination (*11*), we report the brainstem constructs signals for threat probability, fear behaviour, and prediction error from neuronal ‘building blocks’ organized into functional networks. Remapping of neuronal function between cue and outcome periods revealed distinct brainstem network organization for threat probability, fear behaviour, and prediction error signaling.

## Results

Ten rats (4 females) were first shaped to nose poke for food reward. Independent of poke-food contingencies, rats received probabilistic fear discrimination during which three cues predicted unique foot shock probabilities: danger (*p*=1), uncertainty (*p*=0.25) and safety (*p*=0) (Fig. 1A). A 0.25 uncertainty probability was chosen because higher probabilities can produce behavior equivalent to danger (*12*). Rats were implanted with a Neuropixels probe through the brainstem (Fig. 1B) to permit high-density, single-unit recordings from a complete dorsal-ventral axis during discrimination. Fear was calculated with a suppression ratio (see methods), comparing reward seeking rates during baseline and cue periods. Ratio extremes indicated complete suppression of reward seeking (1) versus no suppression (0). Ratios between 0 and 1 indicated intermediate levels of suppression. Rats showed complete discrimination during recording sessions. Suppression of rewarded nose poking was high to danger, intermediate to uncertainty, and low to safety (Fig. 1C; ANOVA main effect of cue, F(_2,142_) = 149.2, *p*=1.26 x 10^-35^; Fig S1). We isolated and held 1812 neurons during 75, 1-hr recording sessions (965 neurons from the 4 females, Table S1). Neurons spanned 21 brainstem regions (*13*) (Fig. 1D), including subregions and neighbouring regions of the superior colliculus, periaqueductal gray, dorsal raphe, and median raphe (Fig. 1E, Fig S2, Table S2).

**Figure 1.**
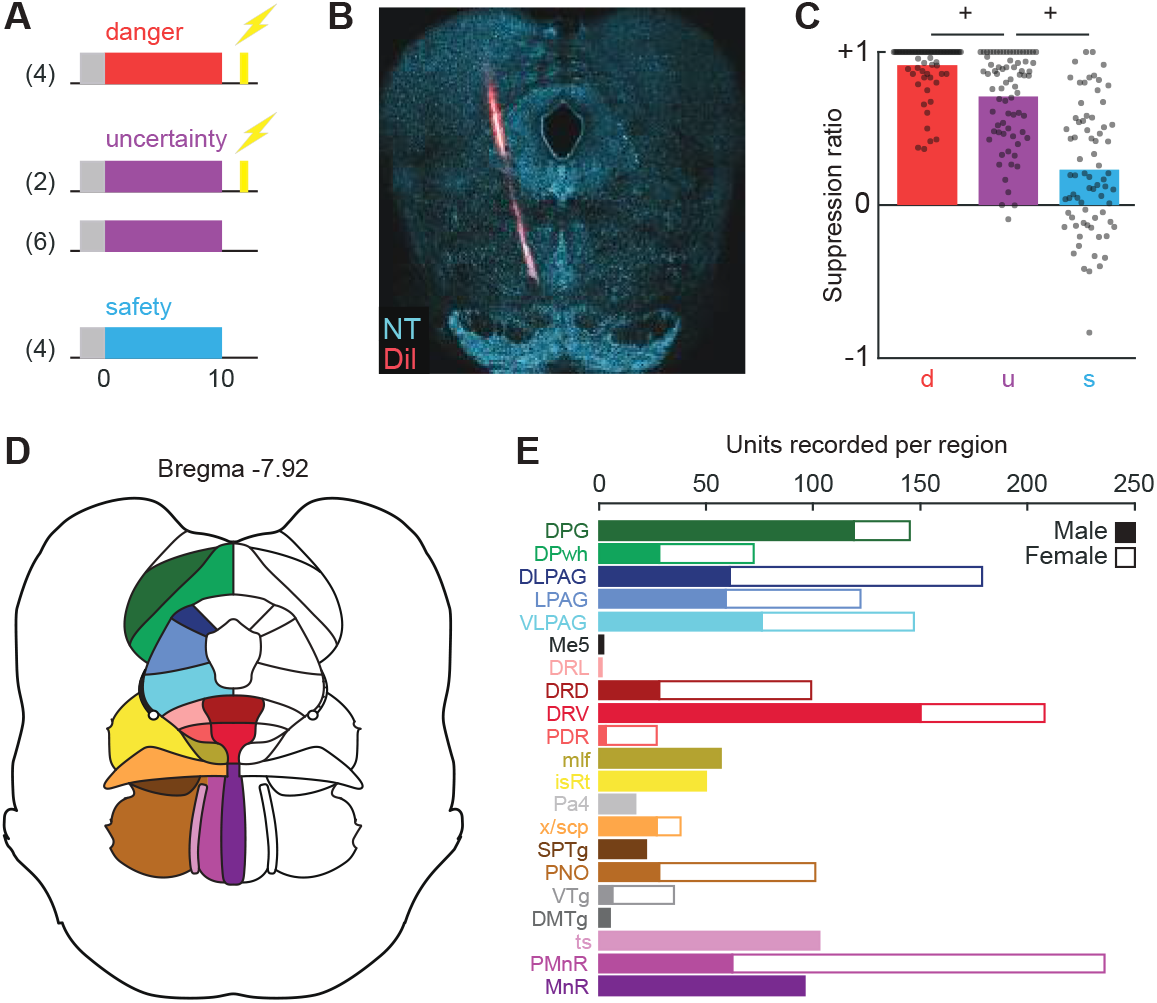
Fear discrimination and Neuropixels implant. (**A**) Probabilistic fear discrimination procedure. (**B**) Representative Neuropixels implant. (**C**) Cue suppression ratios during recording sessions. (**D**) Summary of brainstem regions recorded. (**E**) Number of single units recorded from each brainstem region by sex. ^+^ indicates 95% bootstrap confidence interval does not contain zero.

Brainstem neurons showed marked cue firing that varied in time course, direction, and pattern (Fig. 2A). K-means clustering for mean, single-unit firing in 1-s bins from 2-s prior to 2-s following cue presentation (danger, uncertainty, and safety) revealed neurons could be organized into at least 21 functional clusters; potential building blocks for brainstem construction of threat probability and behaviour. Cluster size varied (min size = 24, max = 219, and median = 83) and consistent firing themes emerged when clusters were visualized (Fig. 2B, Fig S3/S4). Many clusters showed ordered cue firing that differentiated danger, uncertainty, and safety (e.g., k1, k2, k6, k7). Accordingly, principal component analysis (PCA) revealed ordered cue firing (danger > uncertainty > safety) to be the primary low-dimensional feature across all brainstem neurons (PC1, explaining 29.7% of firing variance; Fig. 2C, inset).

**Figure 2.**
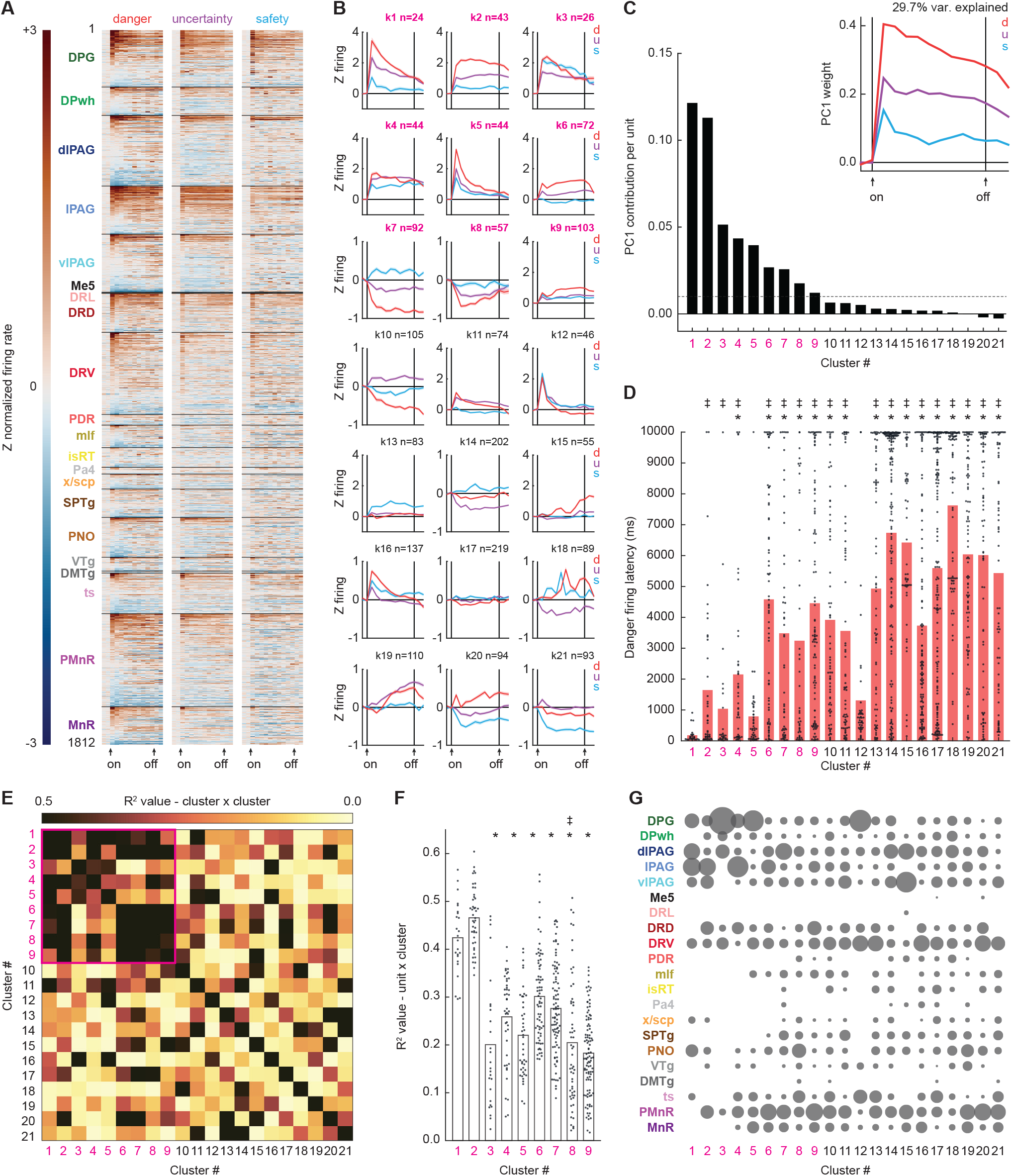
Brainstem cue firing. (**A**) Single-unit firing to danger, uncertainty, and safety cues organized by brain region, dorsal to ventral. (**B**) Mean cluster (k1-k21) firing over cue presentation. (**C**) PC1 for brainstem cue firing (inset) and cluster contribution to PC1. (**D**) Single-unit danger firing latency with cluster means. (**E**) Between-cluster cue firing correlations (data from B). Cue subnetwork outlined in magenta. (**F**) Single unit x cluster firing correlations for the cue subnetwork, with cluster means. (**G**) Proportion of each cluster found in each brainstem region. *Significance of independent samples t-test, Bonferroni corrected. ^‡^Significance of Levene’s test for equality of variance, Bonferroni corrected.

We used an iterative, PCA shuffle analysis to determine the magnitude of each cluster’s PC1 contribution. Cue firing for the neurons comprising a specific cluster (e.g., k1) was shuffled, while cue firing for the neurons of all other clusters was left intact (e.g., k2-k21). Shuffling and PCA were performed 1000 times per cluster. The change in % explained firing variance from the complete data (29.7%) to the shuffled data was calculated and averaged across the 1000 iterations for each specific cluster [PC1 complete – mean (PC1 k1 shuffled)]. PC1 contribution per unit was obtained by dividing the final value by the number of neurons in the cluster. Clusters contributing more greatly to PC1 have higher values.

PC1 firing information largely originated from nine clusters (k1-k9; Fig. 2C, Fig S5) composed of a minority of neurons (505/1812, 27.9%). Clusters k1-k9 showed shorter firing latencies to danger onset, and across all clusters, danger firing latency negatively correlated with the magnitude of PC1 contribution (R^2^ = 0.41, *p*=0.0017; Fig. 2D). Clusters k1-k9 further separated themselves based on cluster-cluster cue firing correlations (Fig. 2E. Fig S5), forming a functional subnetwork within the larger brainstem network. K1 & k2 neurons showed indicators of a subnetwork hub: having the greatest PC1 contributions, and shorter danger firing latencies (Fig. 2D). Further, k1 & k2 single-unit firing correlated most strongly with mean cue firing of their fellow subnetwork clusters (Fig. 2F, Fig S5). Neurons from each cluster were observed in at least six brainstem regions (Fig. 2G). Subnetwork neurons – including k1 & k2 hub neurons – were particularly concentrated in the deep layer of the superior colliculus and subdivisions of the periaqueductal gray.

Ordered cue firing is the predominant brainstem feature. Yet, ordered cue firing could equally reflect fear behaviour or threat probability. Cue firing reflecting behaviour should scale to the level of suppression, invariant of shock probability. Conversely, cue firing reflecting threat probability should linearly scale with shock probability (0.0, 0.25, and 1.0), invariant of the level of suppression. Linear regression revealed unique fear behaviour and threat probability signaling across the 21 clusters (Fig. 3A). Clusters showed considerable temporal variation in signaling prior to and following cue presentation, characterized by greatest signaling at onset (e.g., k5), offset (e.g., k15), sustained over cue presentation (e.g., k10), or even U-shaped signaling peaking mid cue (e.g., k6).

**Figure 3.**
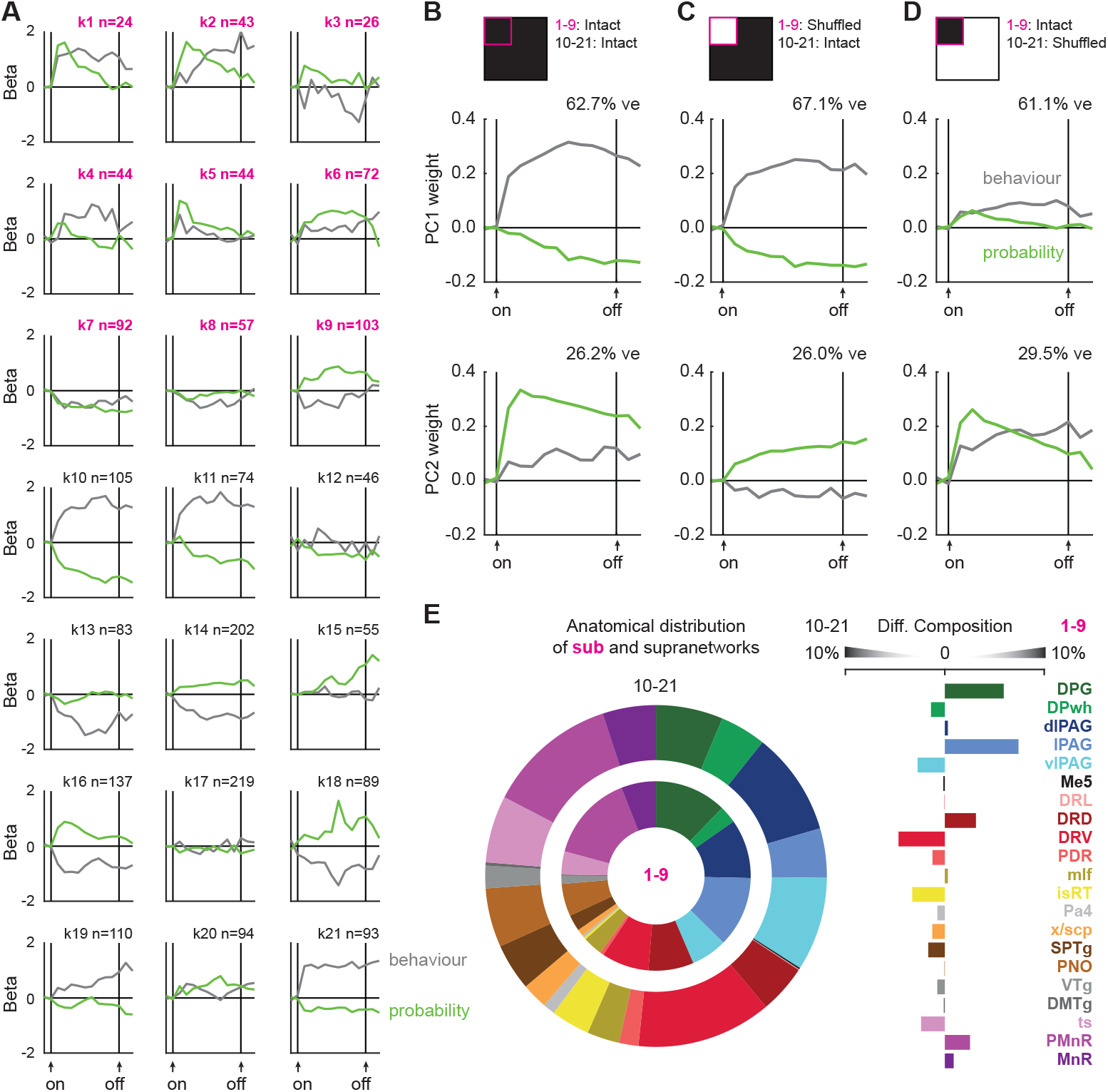
Brainstem threat and behaviour signaling. (**A**) Mean cluster (k1-k21) beta weights for threat probability and behaviour over cue presentation. (**B**) Principal components for cluster beta weights (top; data from A), resulting PC1 (middle), and PC2 (bottom). (**C**) Principal components for cluster beta weights with cue subnetwork shuffled (top; data from A), resulting PC1 (middle), and PC2 (bottom). (**D**) Principal components for cluster beta weights with cue supranetwork shuffled (top; data from A), resulting PC1 (middle), and PC2 (bottom). (**E**). Proportion of cue subnetwork and supranetwork single units by brain region (left), and differential composition of subnetwork and supranetwork by brain region (right).

To reveal low-dimensional signaling features across all clusters (and therefore across the brainstem), we performed PCA on fear behaviour and threat probability beta coefficients prior to and following cue presentation (Fig. 3B, top). PC1 reflected a sustained behaviour signal that peaked mid cue presentation, with lesser and opposing threat probability signaling (62.7% of signaling variance; Fig. 3B, middle). PC2 reflected a sustained threat probability signal that peaked during early cue presentation, with lesser behaviour signaling (26.12% of signaling variance; Fig. 3B, bottom, Fig S5). Thus, stable signals for fear behaviour and threat probability emerged from disparate, temporal signals across clusters.

To reveal network-specific contributions to low-dimensional signals, we iteratively shuffled or ‘lesioned’ cluster firing for one network (e.g., subnetwork clusters k1-k9), while leaving the remaining clusters intact (e.g., supranetwork clusters k10-k21). After network-specific firing shuffling, we performed linear regression for each cluster, then performed PCA for behaviour and threat probability beta coefficients across all clusters. Comparing fully intact signaling (Fig. 3B), to signaling observed with the subnetwork lesioned (Fig. 3C), versus the supranetwork lesioned (Fig. 3D), allowed us to determine the relative contributions of each brainstem network to behaviour and threat probability signaling.

Sustained threat probability signaling depended more on the cue subnetwork, while sustained fear behaviour signaling depended more on the cue supranetwork (Fig. 3C and D). Lesioning the subnetwork left sustained behaviour signaling largely intact (Fig. 3C, PC1, middle), but reduced threat probability signaling, particularly at cue onset (Fig. 3C, PC2, bottom). By contrast, lesioning the supranetwork diminished sustained behaviour signaling (Fig. 3D, PC1, middle), and shifted sustained threat probability signaling to dynamic, probability-to-behaviour signaling (Fig. 3D, PC2, bottom). Neurons contributing to the subnetwork and supranetwork were distributed throughout the brainstem. Subtle anatomical biases were only apparent for the cue subnetwork (Fig. 3E), with the deep layer of the superior colliculus and the lateral subdivision of the periaqueductal gray (also sources of hub neurons) contributing more to the cue subnetwork. Collectively, these findings reveal the brainstem is composed of diverse, neuronal building blocks whose specific cue firing patterns (Fig. 2A) carry unique temporal information about threat probability and fear behaviour (Fig. 3A). A sustained fear behaviour signal is observed across all brainstem neurons (Fig. 3C, middle) while a sustained threat probability signal (Fig. 3D, bottom) is constructed by functional subnetwork.

Fear behaviour and threat probability signals are shaped by prediction errors generated following surprising shock delivery and omission. To capture prediction error-related firing, we focused on the 10 s following shock offset. Brainstem neurons showed marked and varied firing changes, particularly following ‘surprising’ shock on uncertainty trials (Fig. 4A). K-means clustering for mean post-shock firing (4 trial types: danger, uncertainty shock, uncertainty omission, and safety) revealed brainstem neurons could be organized into at least 11 functional clusters (min size = 46, max = 322, and median = 169; Fig. 4B, Fig S6). PCA for mean post-shock firing of all 1812 brainstem neurons revealed positive prediction error, on top of shock responding, to be the primary low-dimensional feature (PC1 explains 17.7% of firing variance, Fig. 4C, Fig S7). The prediction error is ‘positive’ because the PC1 weight for surprising shock delivery (on uncertainty trials) exceeded the PC1 weight for predicted shock (on danger trials). The prediction error is not ‘signed’ because it did not include an opposing, negative error – greater firing decreases to surprising shock omission (on uncertainty trials) versus predicted omission (on safety trials).

**Figure 4.**
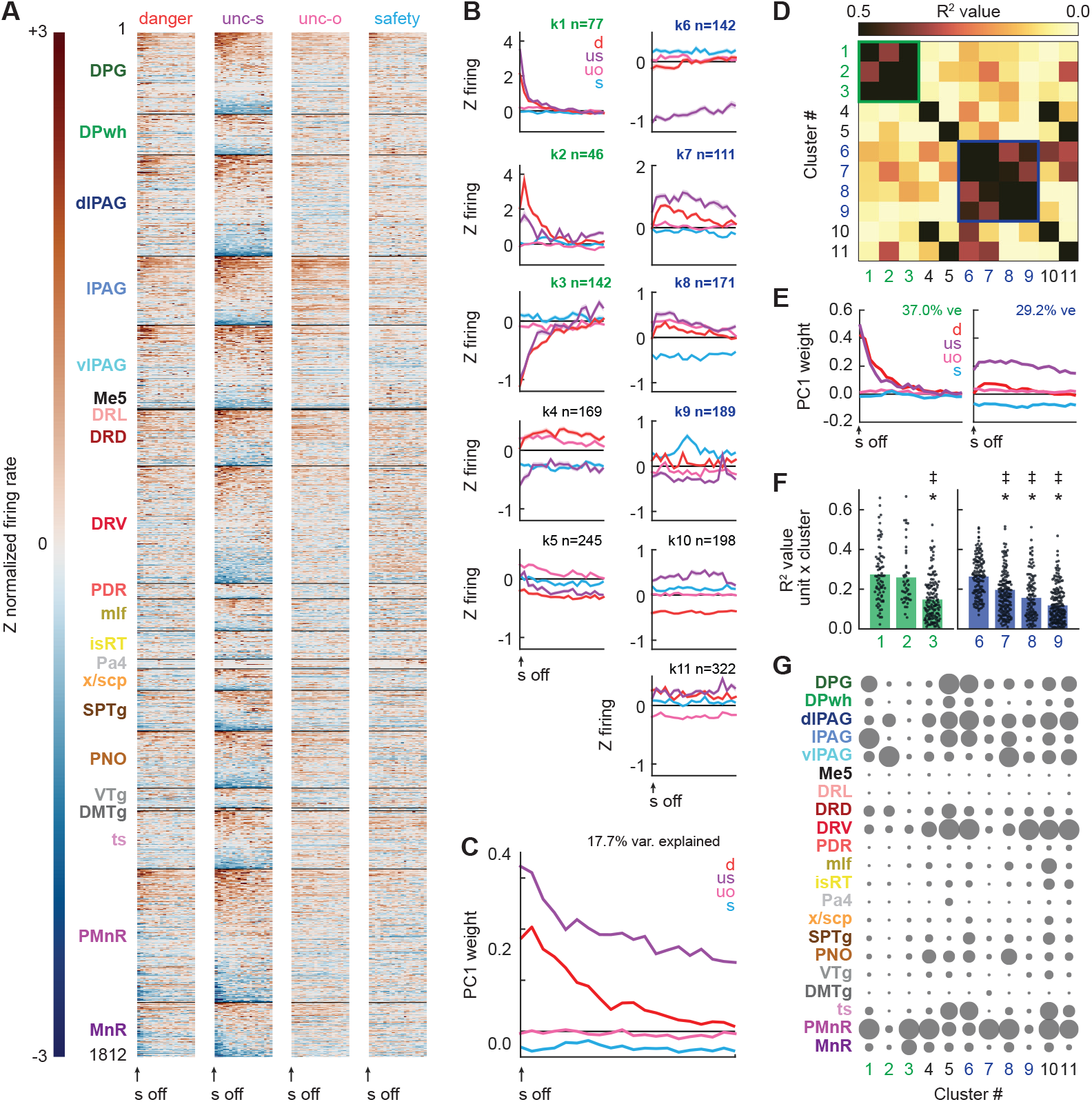
Brainstem outcome firing. (**A**) Single-unit firing following danger, uncertainty shock, uncertainty omission, and safety organized by brain region, dorsal to ventral. (**B**) Mean cluster (k1-k11) firing following shock. (**C**) PC1 for brainstem outcome firing. (**D**) Between-cluster cue firing correlations (data from B). Phasic outcome network outlined in green, tonic outcome network outlined in blue. (**E**) PC1 for phasic outcome network firing (left), and tonic outcome network firing (right). (**F**) Single unit x cluster firing correlations for the phasic outcome network (left, green), and tonic outcome network (right, blue). (**G**) Proportion of each cluster found in each brainstem region. *Significance of independent samples t-test, Bonferroni corrected. ^‡^Significance of Levene’s test for equality of variance, Bonferroni corrected.

Unlike the cue period, the temporal pattern of shock firing organized outcome clusters. K1-k3 neurons showed phasic firing changes following shock. K1 neurons showed greatest, phasic firing increases to surprising shock, and k2 neurons greatest, phasic firing increases to predicted shock, while k3 neurons phasically suppressed firing following shock irrespective of trial type (Fig. 4B, left column). Cluster-cluster firing correlations revealed k1-k3 formed a phasic outcome network (Fig. 4D, Fig S7). K6-k9 neurons showed sustained firing changes following surprising shock (Fig. 4B, right column). K6 neurons selectively inhibited firing to surprising shock, while k7 & k8 neurons showed sustained firing increases that were maximal to surprising shock. Cluster-cluster firing correlations revealed k6-k9 formed a tonic outcome network (Fig. 4D). Network-specific PCA for post-shock firing revealed equivalent and phasic shock firing on danger and uncertainty trials to be the primary low-dimensional feature of the phasic outcome network (explaining 37.0% of firing variance; Fig. 4E, left). PCA further revealed selective firing to surprising shock, and opposing firing to safety, to be the primary low-dimensional feature of the tonic outcome network (explaining 29.2% of firing variance; Fig. 4E, right).

Within the phasic outcome network, k1 & k2 single-unit firing was better correlated with population firing of their fellow network clusters than were k3 neurons (Fig. 4F, left). It is problematic to describe k1 & k2 neurons as hubs, given that the total network contains only 3 clusters. By contrast, k6 neurons were a hub for the tonic outcome network. K6 neuron firing, the cluster strongly decreasing firing to surprising shock, correlated most strongly with population firing of their fellow tonic outcome clusters (Fig. 4F, right). Neurons comprising the phasic and tonic outcome networks differed somewhat in their anatomical distribution (Fig. 4G). Phasic outcome neurons were more common at axis extremes: subregions of the periaqueductal gray, dorsal raphe, and paramedian raphe. Tonic outcome neurons were anatomically diffuse. This distribution was most striking for k6 hub neurons which were evenly distributed throughout the brainstem.

We turned to linear regression in order to distinguish brainstem signals for sensory shock (equating shock firing on danger and uncertainty trials), versus prediction error (differential shock firing on uncertainty trials compared to danger). First, cluster-specific linear regression revealed unique shock and prediction error signals across the 11 outcome clusters (Fig. 5A). For example, k1 neurons transiently signaled both sensory shock and prediction error, while k6 neurons exclusively signaled error. PCA for shock and prediction error beta coefficients across all clusters (therefore all brainstem neurons) revealed opposing sensory shock and prediction error signals to be the primary low-dimensional feature (PC1, 53.7% of signaling variance; Fig. 5B, middle). PC2 reflected dynamic, sensory shock to prediction error signaling (33.4% of signaling variance; Fig. 5B, bottom, Fig S7).

**Figure 5.**
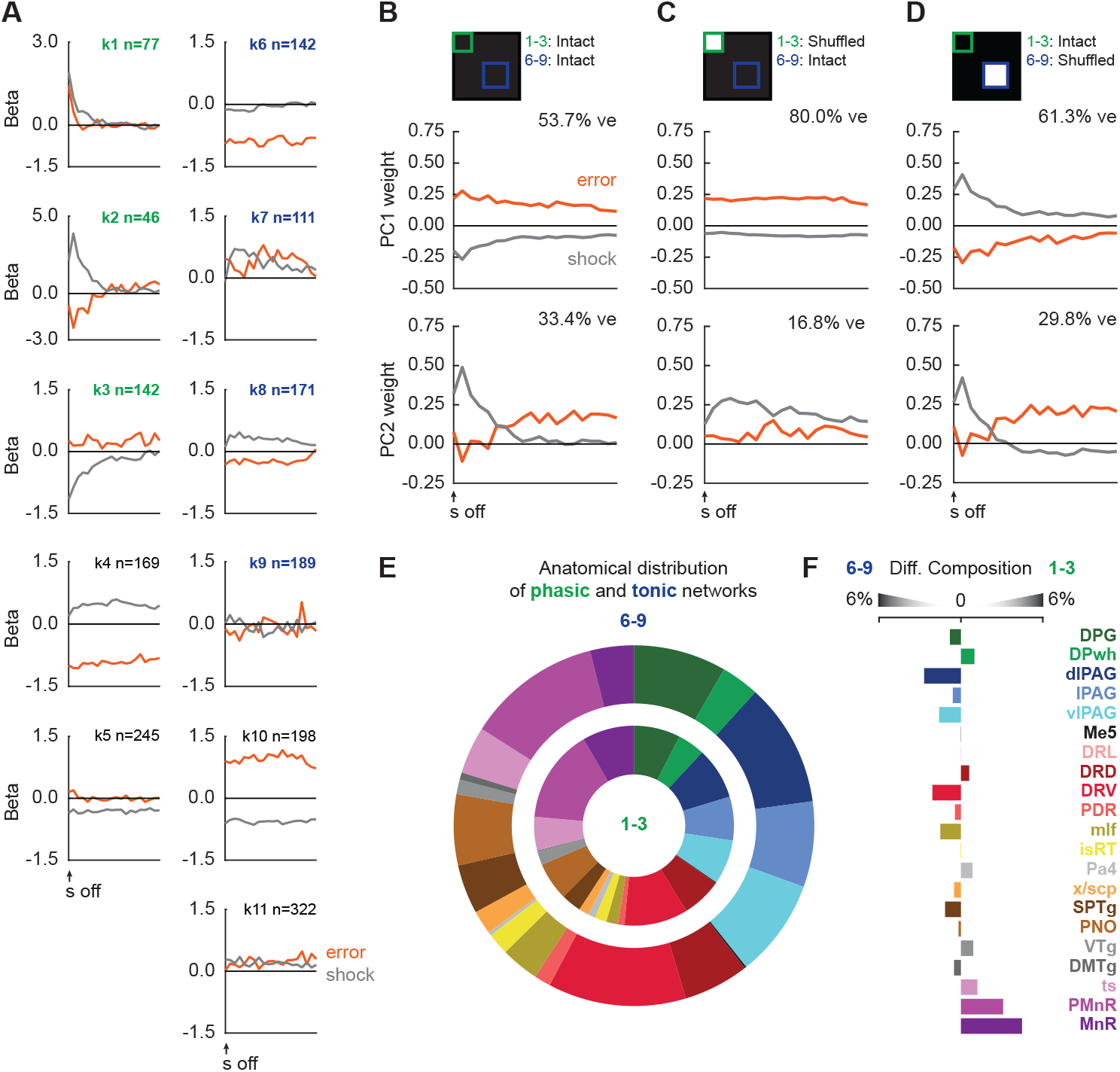
Brainstem shock and prediction error signaling. (**A**) Mean cluster (k1-k11) beta weights for shock and prediction error following shock delivery and omission. (**B**) Principal components for cluster beta weights (top; data from A), resulting PC1 (middle), and PC2 (bottom). (**C**) Principal components for cluster beta weights with phasic outcome network shuffled (top; data from A), resulting PC1 (middle), and PC2 (bottom). (**D**) Principal components for cluster beta weights with tonic outcome network shuffled (top; data from A), resulting PC1 (middle), and PC2 (bottom). (**E**) Proportion of phasic and tonic network single units by brain region (left), and differential composition of tonic and phasic network by brain region (right). (**F**) Relationship between cue cluster and outcome cluster membership across all brainstem neurons.

Rapid sensory shock signaling depended on the phasic outcome network (Fig. 5C). Lesioning the phasic outcome network (using a PCA lesion approach identical to the cue networks) left signaling dominated by sustained prediction error (80.0% of signaling variance; Fig. 5C, middle), while the residual reflected sustained shock (16.8% of signaling variance; Fig. 5C, bottom). By contrast, lesioning the tonic outcome network emphasized opposing, phasic sensory shock and prediction error signaling (61.3% of signaling variance; Fig. 5D, middle). Residual signaling reflected dynamic, sensory shock to prediction error (29.8% of signaling variance; Fig. 5D, bottom), similar to that observed across all brainstem neurons (Fig. 5B, bottom). Thus, a phasic brainstem network signals sensory shock *and* prediction error from a subset of neuronal building blocks transiently responsive following shock. Concurrently, a tonic brainstem network preferentially constructs positive prediction error from a subset of neuronal building blocks sustaining activity following shock. Anatomical biases for phasic and tonic outcome network neurons were nearly absent (Fig. 5E).

The same 1812 neurons constructing threat probability and behaviour during cue presentation, signaled shock and prediction error following shock delivery. We were curious whether there was a relationship between network membership during cue and outcome periods. Recall that the cue subnetwork was composed of 505 neurons (27.9% of all neurons), many fewer neurons than the cue supranetwork (1307/1812, 72.1%). Given this imbalance, the relevant question is if the proportion of phasic (n = 265) and tonic (n = 613) outcome neurons contributing to the cue subnetwork differ from 27.9%. Chi-squared testing found neither the phasic outcome neurons (85/265, 32.1%; χ^2^ = 2.27, *p*=0.13) nor the tonic outcome neurons (159/613, 25.9%; χ^2^ = 1.17, *p*=0.28) differed from the expected proportion of 27.9%. A neuron’s membership to cue networks constructing threat probability and fear behaviour had no influence on it’s membership to outcome networks signaling sensory shock and prediction error.

## Discussion

We set out to reveal brainstem signaling of threat probability versus fear behaviour. Supporting a prevailing view (*1*), we observed functional populations whose firing was better captured by trial-by-trial fluctuations in behaviour, rather than threat probability. The firing independence of these populations and their diffuse anatomical distribution meant continuous brainstem behaviour signaling from cue onset through shock delivery. Opposing the prevailing view, functional populations whose firing was better captured by threat probability were also observed. Continuous brainstem threat probability signaling was also achieved by anatomically diffuse functional populations. However, an organized threat probability signal was uncovered. Brainstem populations showing pronounced differential firing to danger, uncertainty, and safety, plus short-latency danger firing changes, formed a functional subnetwork. Subnetwork ‘hub’ neurons were concentrated in the deep layer of the superior colliculus and periaqueductal gray. Rather than being the exclusive domain of the forebrain, the brainstem constructs threat probability.

We further found that prediction error signaling is fundamental to the brainstem. This is broadly consistent with prior studies which have reported prediction error in the periaqueductal gray (*7, 8, 14*). However, our results paint a more complex picture. First, the brainstem contains two networks operating on different time scales. A phasic outcome network is rapidly engaged following shock, with composing functional populations generally inhibited to shock, or showing selective phasic firing increases to surprising or predicted shock. Populations for surprising shock – positive error – overlap in known centers for prediction error generation (*8*). A tonic outcome network specifically signals positive prediction error. Unique to the tonic outcome network: composing functional populations – even the hub – are highly anatomically distributed. Even more, hub neurons show preferential firing *decreases* to surprising shock. The results suggest there are either multiple brainstem prediction error systems, or there is a single prediction error system in which a tonic signal opens a window of permissibility for phasic error to update threat estimates (*15*).

A caveat to our prediction error findings is that we did not observe population-level or brainstem-level signaling of *negative* error. The intermediate levels of nose poke suppression to the uncertainty cue confirm that shock omission was detected – otherwise uncertainty behavior would have been equivalent to danger. Though because we selected a 25% foot shock probability for the uncertainty cue, shock trials were rarer than omission trials. There is evidence that midbrain dopamine neurons signaling prediction error are additionally sensitive to rare outcomes (*16*). Had we made shock omission rarer than shock delivery (e.g., using a 75% shock probability), neural correlates of negative error might have emerged. Yet given that a 50% shock probability cue can support behavior comparable to a 100% cue (*12*), negative error generated by surprising omission to a 75% shock cue would likely be insufficient to weaken cue-shock associations and reduce fear behaviour.

Viewing the forebrain as *the* source of threat estimation has meant continuous refinement of forebrain threat processing. Cortical subregions are linked to increasingly specific threat functions (*17*). Amygdala threat microcircuits are being mapped in intricate detail (*18*). Brainstem regions contain the building blocks needed to construct threat estimates. This finding necessitates refinement and detail of brainstem threat function on par with its forebrain counterparts. Expanding on prior brainstem work (*19–21*), our results reveal the superior colliculus (*22–24*) and periaqueductal gray (*25–27*) as prominent sources of threat information. Somewhat unexpectedly, we reveal abundant and diverse threat signaling in the paramedian raphe (*28*), a virtually unstudied region adjacent to the serotonin-containing median raphe. Critically, these regions do not function in isolation. Rather, the superior colliculus and periaqueductal gray organize a local brainstem network to signal threat probability.

Perhaps the brainstem signals threat probability, but this signal is trained up by the forebrain. This would be consistent with our findings. Yet, where do forebrain threat estimates originate and once formed, how are threat estimates updated? Prediction error provides a plausible mechanism for forming and updating threat estimates. Preferential responding to surprising aversive events – consistent with positive prediction error – has been reported in many human forebrain regions (*29*). Preferential responding to omission of aversive events – consistent with negative error – has also been observed in human forebrain regions (*30*). However, opposing firing changes to positive and negative error in the same region – a requirement of a *signed* prediction error (*31, 32*) – are more narrowly observed in the human brainstem (*8*). Direct manipulation of error-related activity in the brainstem of rats and mice alters fear behavior (*3, 7, 33, 34*). Available evidence suggests that learned threat estimates originating in the forebrain require prediction error generated in the brainstem. In which case, *de novo* acquisition of a brainstem threat estimate, trained by the forebrain, would require a brainstem-generated prediction error. Also plausible – brainstem-generated prediction error may train and update a brainstem threat estimate, bypassing the forebrain altogether.

Fully revealing the brain basis of threat computation is essential to understanding adaptive and disordered fear. Our finding of widespread and organized brainstem threat signaling calls for abandonment of the historical division of labour view. In its place we must embrace a brain-wide view of threat computation (*36–38*) in which brainstem networks are not limited to organizing fear behaviour but are integral to estimating threat.

## Acknowledgments

We thank Bret Judson and the Boston College Imaging Core for infrastructure/support, Dr. Matthew Gardner and Dr. Geoffrey Schoenbaum for advice and initial designs for 3D head cap printing, Joe Austen for help building and ordering acquisition computers, Richard Pijar and the Boston College Machine Shop for electrical and hardware support and instrument customization, Pavel Kulik and Open Ephys for assistance with the I/O module, Dr. Nicholas Steinmetz, Dr. Matteo Carandini and the UCL Neuropixels course for initial training in the use of silicone probes. Research reported in this publication was supported by the National Institute of Mental Health of the National Institutes of Health under Award Number R01MH117791. The content is solely the responsibility of the authors and does not necessarily represent the official views of the National Institutes of Health. The authors report no competing interests. This work was also supported in part by an Ignite Grant from Boston College.

## Funding

R01-MH117791

## Author contributions

Conceptualization: JS, MM

Methodology: JS, MM

Investigation: JS, MM

Funding acquisition: MM

Writing – original draft: JS, MM

Writing – review and editing: JS, MM

## Competing interests

Authors declare that they have no competing interests.

## Data Availability

Data will be uploaded and available at https://crcns.org/

## Code Availability

All code used for single unit data sorting and analysis will be uploaded and available at https://crcns.org/

## Materials and Methods

### Subjects

Subjects were six male and four female Long-Evans rats, split over two rounds of testing. The first round included three female and two male rats born in the Boston College Animal Care facility, housed with mothers until postnatal day 21 when they were weaned and single housed. The second round included four males and one female, obtained from Charles River weighing 250g-275g on arrival. All were maintained on a 12-hour light-dark cycle (lights on 0600–1800) and were aged between 95 - 140 days old at the time of first recording session. All protocols were approved by the Boston College Animal Care and Use Committee, and all experiments were carried out in accordance with the NIH guidelines regarding the care and use of rats for experimental procedures.

### Behavioural apparatus

Training took place in individual sound-attenuated enclosures that each housed a behaviour chamber with aluminum front and back walls, clear acrylic sides and top, and a metal grid floor. Each grid floor bar was electrically connected to an aversive shock generator (Med Associates, St. Albans, VT) through a device that ensured the floor was always grounded apart from during shock delivery. A single food cup and central nose poke opening equipped with infrared photocells were present on one wall. Auditory stimuli were presented through two speakers mounted on the enclosure ceiling. Auditory cues were 10s in duration and consisted of repeating motifs of a broadband click, phaser, or trumpet, which previous studies have found to be discriminable and equally salient. Testing took place in an identical chamber, but was equipped with a custom plastic food cup, plastic front and back walls, and multi-axis counterbalanced lever arm (Instech Laboratories, MCLA) with plastic tubing that held the recording cable and entered the chamber via a custom plastic top.

### Nose poke acquisition

Rats were food restricted to 85% of their free-feeding body weight, with ad-libitum access to water. After pre-exposure to pellets (Bio-Serv, Flemington, NJ) in their home cages for two days, rats were shaped to nose poke for pellet in the experimental chamber. During the first session, the nose poke port was removed, and rats were issued one pellet every 60 seconds for 30 minutes. In the next session, the port was reinserted, and poking was reinforced on a fixed ratio 1 schedule in which one nose poke yielded one pellet until they reached ∼50 nose pokes or 30min. Nose poking was then reinforced on a variable interval 30-second (VI-30) schedule for one session, then a VI-60 schedule for the next four sessions. The VI-60 reinforcement schedule was utilized during subsequent fear discrimination and was independent of auditory cue and shock presentation.

### Fear discrimination

Rats received twelve sessions of Pavlovian fear discrimination prior to Neuropixels implant. Each 54-min session consisted of a five-minute warm up period in the chamber followed by 16 cue presentation trials. Each auditory cue predicted a unique shock probability (0.5 mA, 0.5 s): danger, *p*=1.00; uncertainty, *p*=0.25; and safety, *p*=0.00. Shock was administered two seconds following the termination of the cue on danger and uncertainty-shock trials. A single session consisted of 4 danger, 2 uncertainty-shock, 6 uncertainty-no shock, and 4 safety trials with a mean inter-trial interval of 3 min. Trial order was randomly determined by the behavioural program and differed for each rat, every session. The physical identities of the auditory cues were counterbalanced across individuals. Following recovery from surgery, rats received one VI-60 session to habituate to being connected to the recording cable. Rats then received between 1 and 10 discrimination sessions during which single-unit activity was recorded.

### Calculating suppression ratio

Time stamps for cue presentations, shock delivery and nose pokes (photobeam break) were automatically recorded by the Med Associates program. Baseline nose poke rate was calculated for each trial by counting the number of pokes during the 20-s pre-cue period and multiplying by 3. Cue nose poke rate was calculated for each trial by counting the number of pokes during the 10-s cue period and multiplying by 6. Nose poke suppression was calculated as a ratio: (baseline poke rate – cue poke rate) / (baseline poke rate + cue poke rate). A suppression ratio of ‘1’ indicated complete suppression of nose poking during cue presentation relative to baseline. A suppression ratio of indicated ‘0’ indicates equivalent nose poke rates during baseline and cue presentation. Gradations in suppression ratio between 1 and 0 indicated intermediate levels of nose poke suppression during cue presentation relative to baseline.

### Surgery

Following the 12^th^ discrimination session rats were returned to ad-libitum food access and underwent stereotaxic surgery performed under isoflurane anesthesia (1-5% in oxygen). Four screws were screwed into the skull around the target cap area to aid adhesion of the cap, and the skull was also scored in a crosshatch pattern. A craniotomy with a 1.4 mm diameter was carried out, and the underlying dura fully removed to expose the cortex. Immediately prior to implant the probe was painted with Dil to later identify histology tracks (ThermoFisher, V22886). To maximize recording regions, each implant was aimed at coordinates −8.00 AP, −2.80 ML, −7 to −7.5 DV, with a 15⁰ angle. Each Neuropixels probe (1.0 probe) and head stage were secured in a pre-prepared custom head cap. The cap was held and slowly lowered during implant using a modified stereotaxic arm until the max DV was reached, or until the cap contacted the skull. The craniotomy was sealed using silicone gel (Dow DOWSIL 3-4680). Once the cap was in place, the ground wire was wrapped around the two screws positioned laterally to the cap to ground the probe. Vacuum sealing grease (Dow Corning) was applied around the base of the cap to fill any space between the cap and the skull and protect the probe. Caps were cemented into place using orthodontic resin (cc 22-05-98, Pearson Dental Supply) and the head cap lid secured in place on the head cap. Rats were given one week to recover with prophylactic antibiotic treatment (cephalexin, Henry Schein Medical) prior to data acquisition and received carprofen (5mg/kg) for postoperative analgesia.

### Data acquisition

Neural data were recorded using OpenEphys with the Neuropixels PXI plugin running on an acquisition computer connected to the PXI chassis (PXIe-1071) containing the Neuropixels base station. Behaviour events were controlled and recorded by a separate computer running Med Associates software. To get behaviour timestamps, signals were sent from Med Associates to the NIDAQmx OpenEphys plugin, via Med Associates TTL adapter boxes (SG-231) plugged into a connector block (National Instruments, BNC 2110) connected to an I/O module (PXI-6363) in the PXI chassis. During recording sessions, the cable was first connected to the head stage and the head stage lid fixed in place, then the recording channels and reference for that session and subject were selected. To maximize acquisition of neurons from the midbrain region, the channels selected were either the lowest bank of 384 channels, or channels 193-575, used in a double alternating order across sessions and counterbalanced across subjects. The external reference was selected unless that proved ineffective in which case the tip reference of the probe was used instead. After this the doors to the chamber were closed and the fear discrimination and recording session started. Sessions were only included for analysis if the probe signal was maintained throughout all 16 trials, if the signal was lost for any reason that session was discarded. Subjects were recorded from daily up to either ten total recording sessions, or until data was no longer able to be acquired from a subject.

### Probe retrieval

Following recording sessions rats were placed back into the stereotaxic frame under isoflurane anesthesia. The head cap lid was removed, the ground wire cut, and the head stage disconnected and removed. The cement securing the probe holder in place was scraped away with a scalpel blade and the holder slowly pulled up and out of the cap. The probe was then rinsed and soaked in DI water, followed by a soak in a tergazyme solution before a final rinse with DI water before and if still functional after explant safely stored for re-implant.

### Histology

Once the probe had been explanted, the rat was removed from the frame and deeply anesthetized using isoflurane before being perfused intracardially with 0.9% biological saline and 4% paraformaldehyde in a 0.2M potassium phosphate buffered solution. Brains were extracted and fixed in a 10% formalin solution for 24hrs, then stored in 10% sucrose/formalin. Brains were sliced with a microtome into forty micrometer sections (from approximately Bregma −6.5 to −9, to ensure the full extent of the probe tracks could be identified). The tissue was rinsed, incubated in NeuoroTrace (ThermoFisher, N21479), rinsed again, and then mounted prior to imaging within a week of processing (Axio Imager, Z2, Zeiss) to locate probe placement using the visible Dil tracks and NeuroTrace. Neuron locations were established by identifying the 3D location of the tip of the probe relative to the Allen Atlas, as well as the location in which the probe entered the brain (not including the cortex) and the vector of implant calculated. These were also checked against expected electrophysiology patterns (regions of expected low and high activity) for location accuracy.

### Neuron sorting

See supplement X-X for full description. Data were automatically spike sorted using Kilosort 2 (*39*) or 2.5 (https://github.com/MouseLand/Kilosort). Clusters identified by Kilosort were manually curated in Phy (https://github.com/cortex-lab/phy). To assess if activity reflected single-unit activity, inter-spike interval, waveform shape, firing rates, activity change across channels, were all examined. Neurons were also assessed for potential merges with similar nearby clusters, and for potential splitting out of noise/other neurons. Neurons were only kept for analysis if the pattern of activity was confidently identified as neuron activity, and not noise or multi-unit activity. Accepted neurons were finally screened using Matlab and only kept if consistently recorded throughout all trials in the session recorded in, any neurons that showed clear drop offs or loss of recordings were discarded. 52% of neurons accepted in Phy passed Matlab screening.

### Analysis

Matlab was used to extract, collate, and analyze the single-unit data and behaviour timestamp events. Fear was measured by suppression of rewarded nose poking (baseline poke rate – cue poke rate)/(baseline poke rate + cue poke rate). Perceptually uniform color maps were used to prevent visual distortion of the data (*40*). K-means clustering was performed by systematically varying the number of clusters and examining the output for over/under clustering. Single-unit and population firing analyses utilized k-means clustering, principal components analysis, linear regression combined with iterative shuffling. Complete descriptions of single-unit analyses provided in supplement.

---

**Figure S1.**
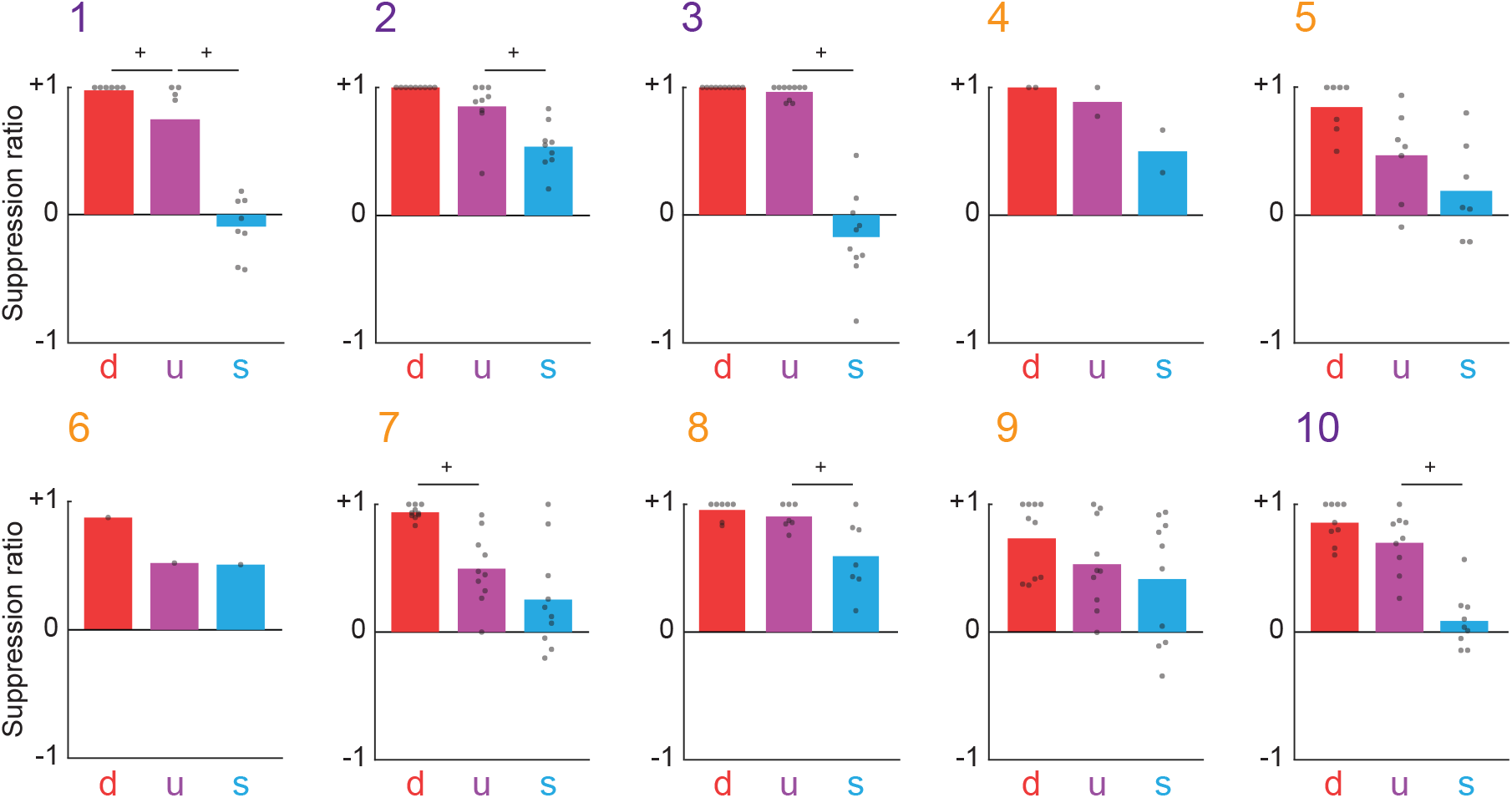
Discrimination plot spreads for each subject. Female (1-3, 10) male (4-9).^+^95% bootstrap confidence intervals do not contain zero.

**Figure S2.**
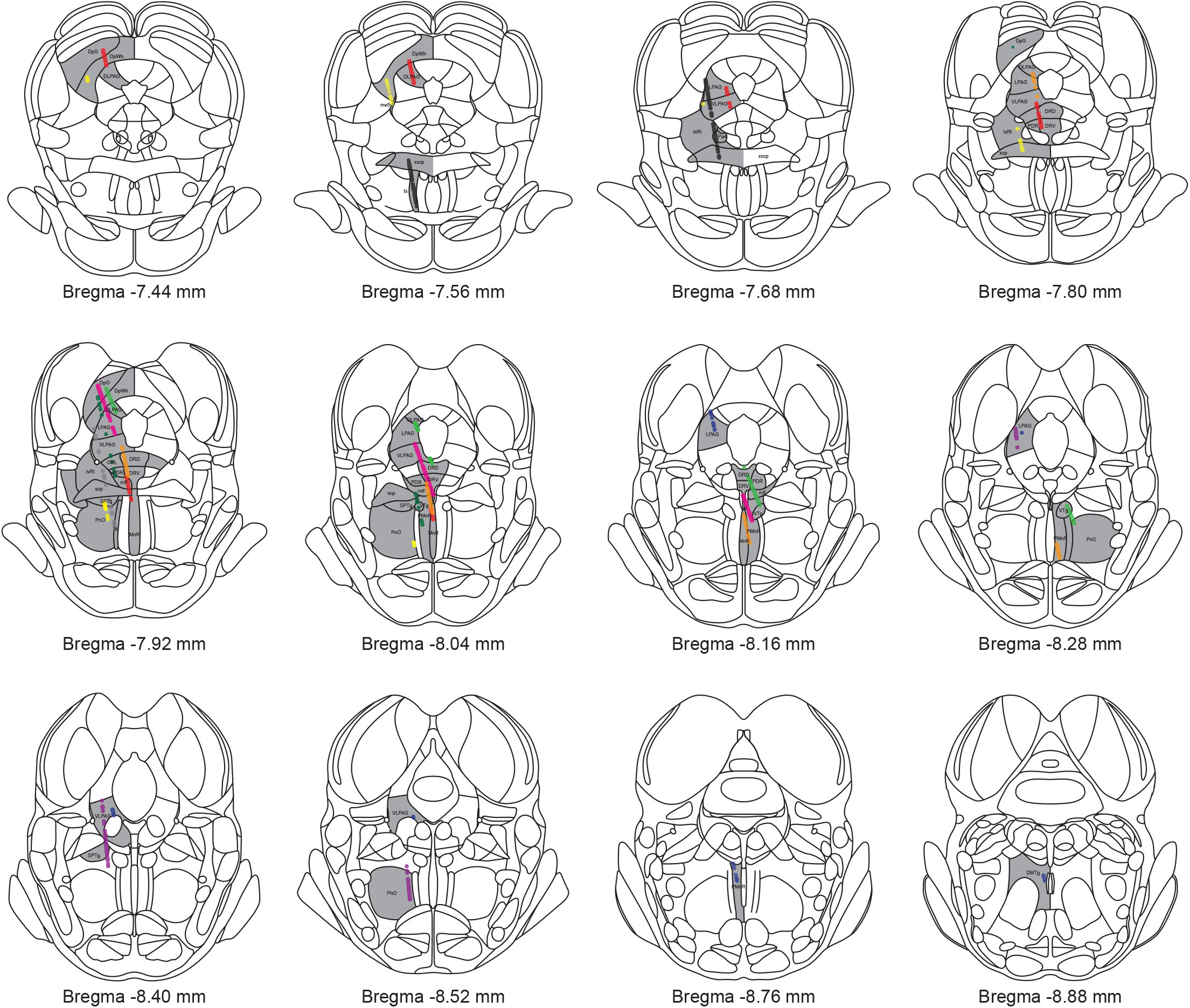
Location of single units for each implant. Subject 1: red, 2: orange, 3: purple, 4: dark green, 5: dark blue, 6: yellow,7: gray, 8: black, 9: pink, 10: light green. Brain regions: deep gray layer superior colliculus (DPG). Deep white layer superior colliculus (DpWh). Dorsolateral periaqueductal gray (DLPAG). Lateral periaqueductal gray (LPAG). Ventrolateral periaqueductal gray (VLPAG). Mesencephalic trigeminal tract (me5). Dorsal raphe nucleus, lateral part (DRL). Dorsal raphe nuclues, dorsal part (DRD). Dorsal raphe nucleus ventral part (DRV). Posterodorsal raphe nucleus (PDR). Medial longitudinal fasciculus (mlf). Isthmic reticular formation (isRt). Paratrochlear nucleus (pa4). Decussation/superior cerebellar peduncle (x/scp). Subpenduncular tegmental nucleus (SPTg). Pontine reticular nucleus, oral part (PnO). Ventral tegmental nucleus (VTg). Dorsomedial tegmental area (DMTg). Tectospinal tract (ts). Paramedian raphe nucleus (PMnR). Median raphe nuclues (MnR).

**Figure S3.**
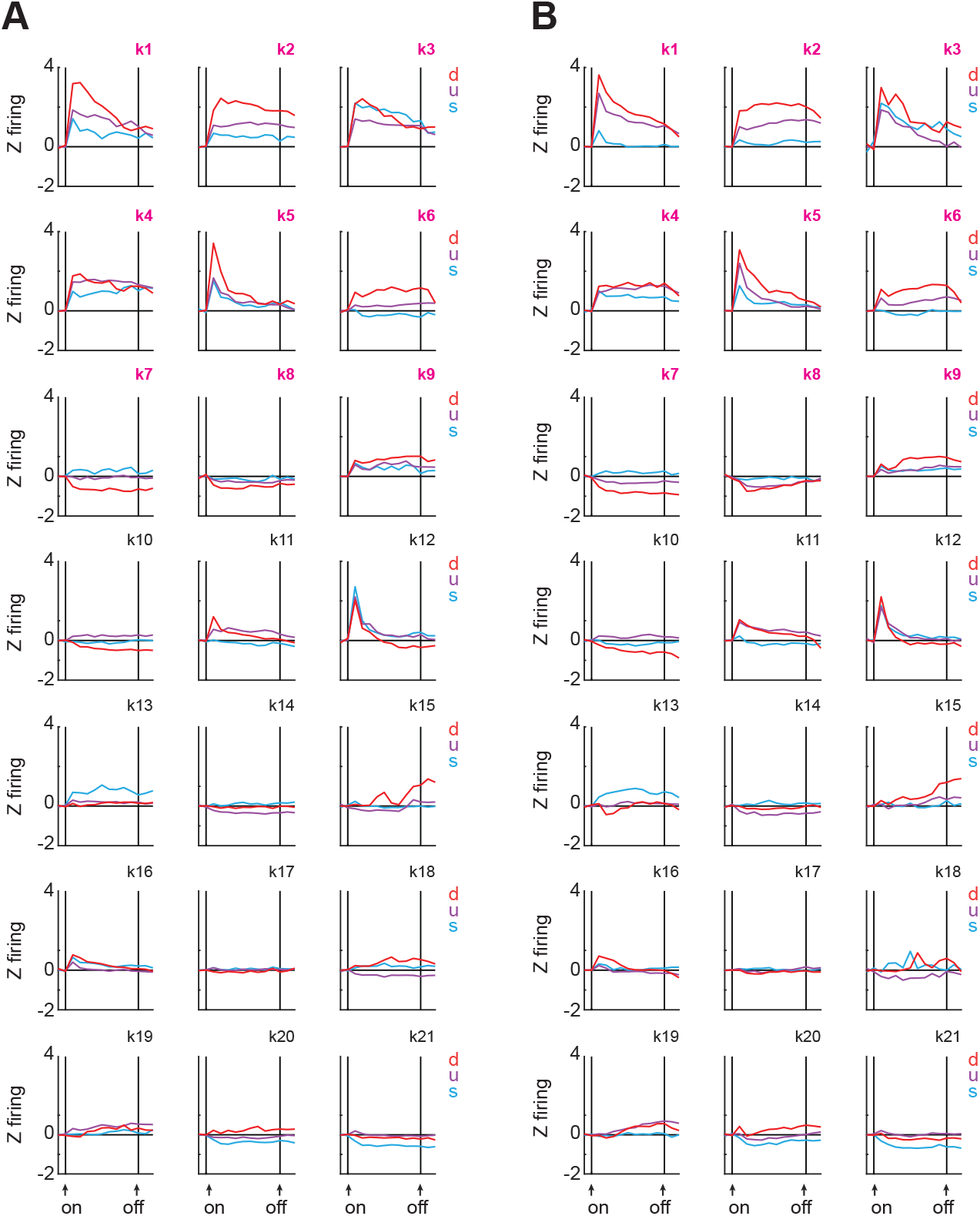
Population firing of 21 clusters during the cue period for female (**A**) and male (**B**) single units.

**Figure S4.**
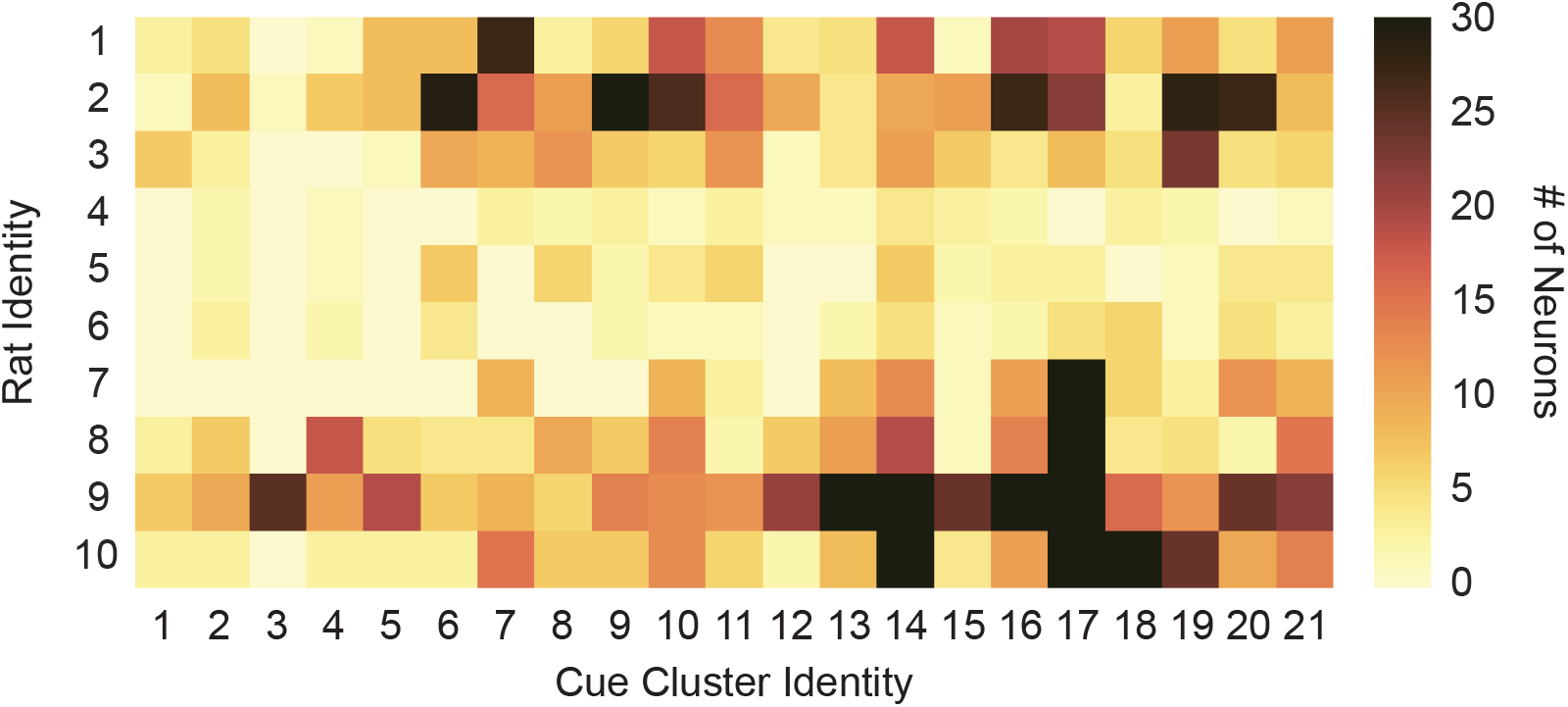
Rat identity (row) x cue cluster identity (column), is plottted for all 1812 neurons.

**Figure S5.**
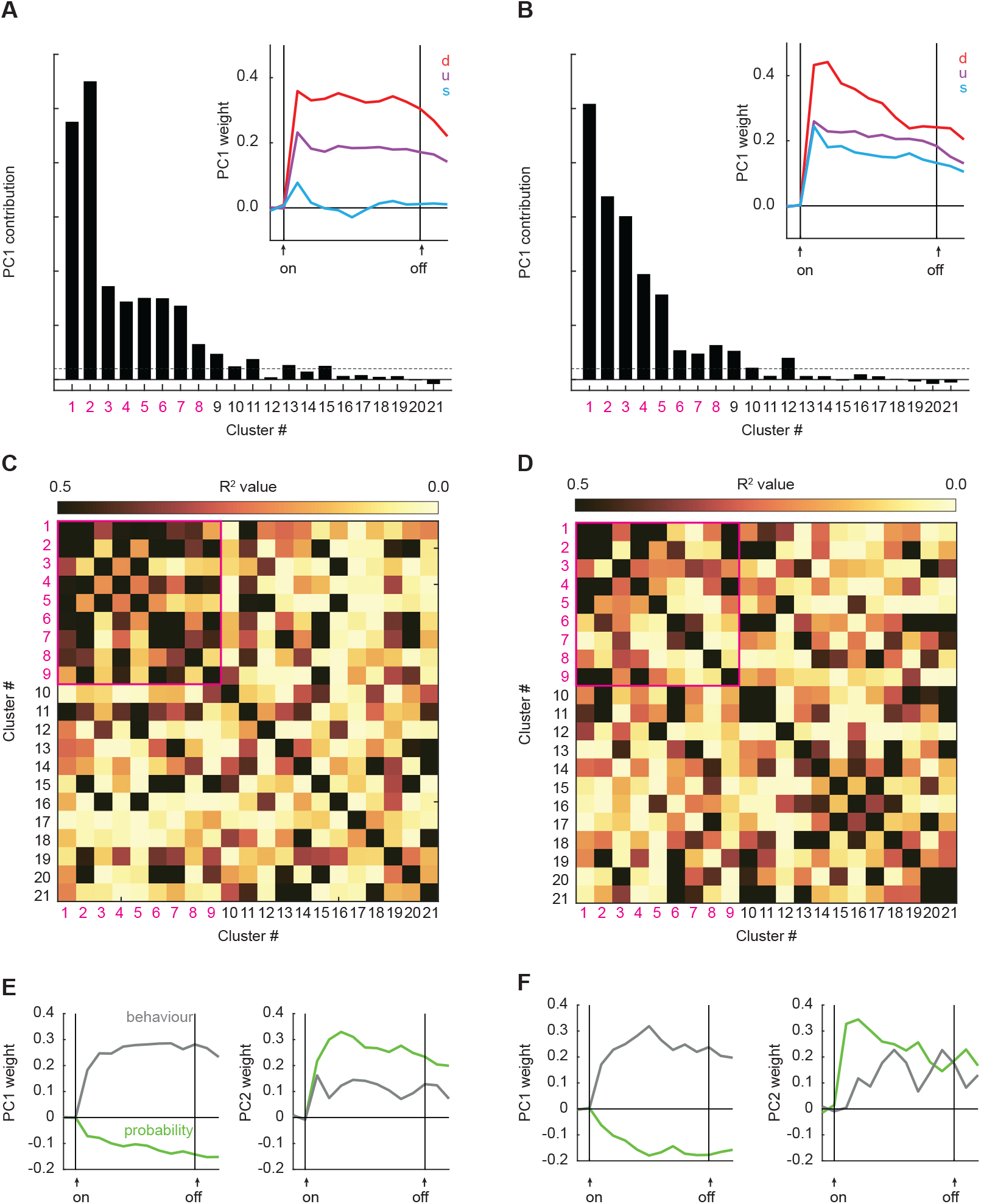
Contributions of PC1 to each cluster and PC1 line plot for female (**A**) and male (**B**) units. Between cluster firing correlations for female (**C**) and male (**D**) units. PCA on regression betas for female (**E**) and male (**F**) single units.

**Figure S6.**
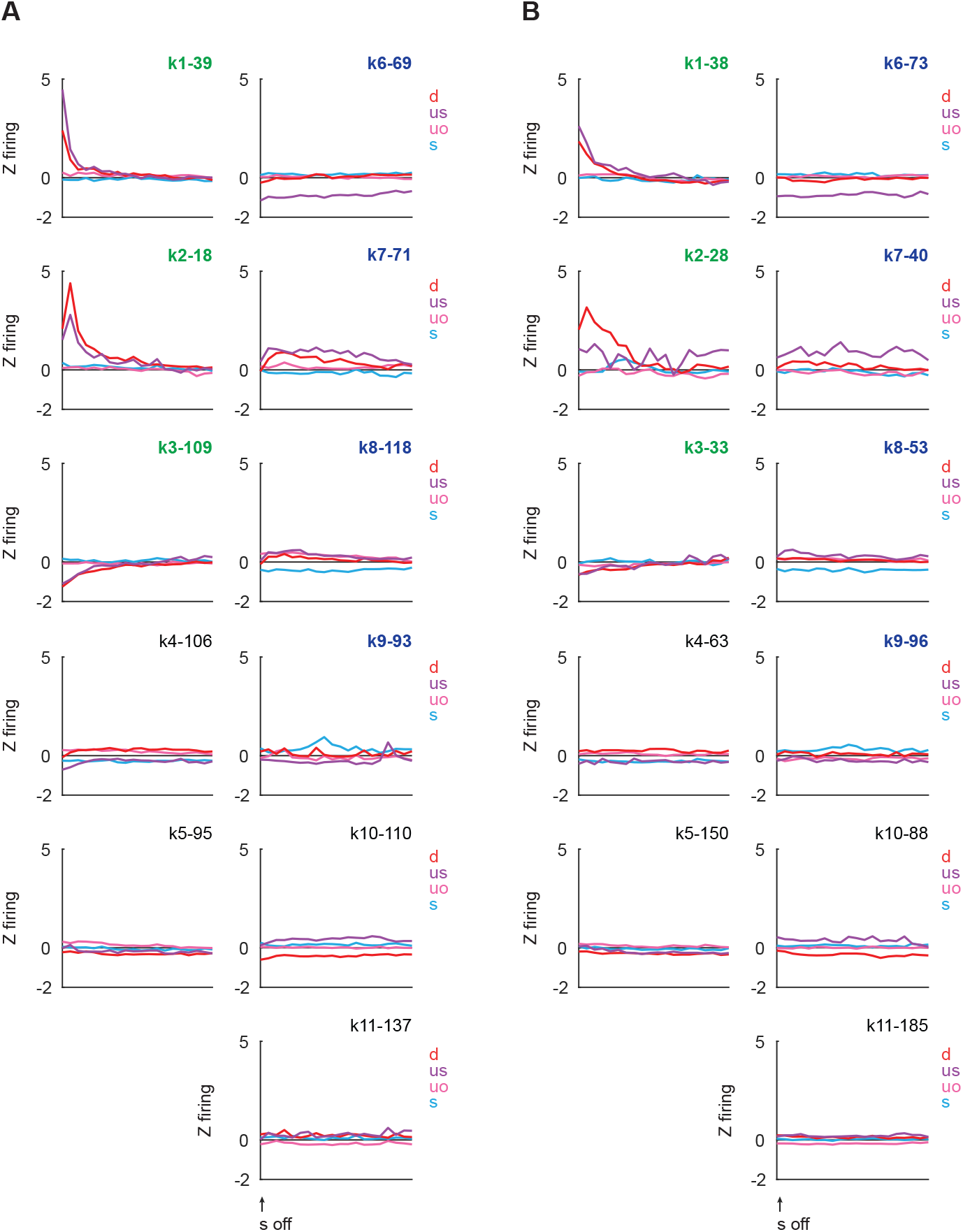
Population firing of 11 clusters during the outcome period for female (**A**) and male (**B**) single-units.

**Figure S7.**
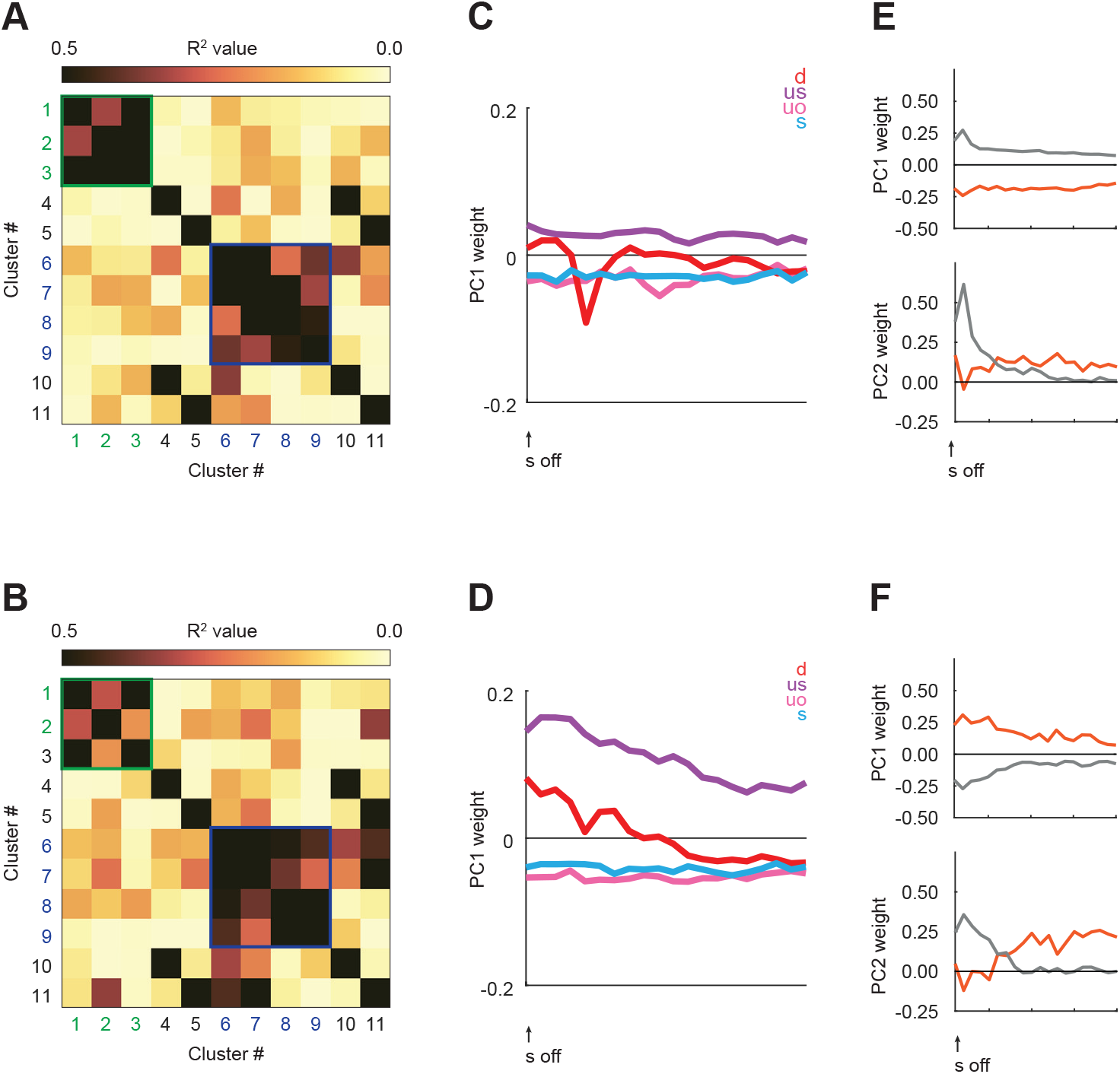
Between cluster firing correlations for female (**A**) and male (**B**) single-units. PCA line plot for female (**C**) and male (**D**) firing. PCA on regression betas for female (**E**) and male (**F**) single-units.

## Head cap preparation

Head cap designs were created using FreeCAD and Tinkercad then 3D printed using a Form2 printer with clear resin (RS-F2-GPCL-04). Prints were washed and rinsed with 90% ethanol before drying for at least 48hrs. The head cap consists of a base section, lid, and probe mount. The probe mount slots into the front section of the cap and the head stage fits into the back section. Neuropixels probes were glued to the probe mount using a small amount of superglue on the back of the probe base. Stainless steel ground wire was soldered to the ground and reference pads located at the top of the wings of the probe. The head stage was attached to the flex cable before the probe mount/probe was carefully inserted into the front section of the head cap, with the ground wire inserted through a small hole in the bottom of the cap. The head stage sits in the back part of the cap with the flex cable forming an S-shape inside the cap. The probe mount and head stage are then fixed in place with small dots of dental cement on the edges, being careful not to prevent the lid from fitting over the base.

The lid is secured with fine adjustment nylon cable ties (McMaster Carr, 6614K11). A separate lid was designed and printed for recording sessions and attached to plastic cable shielding. The recording cable could then run through the shielding and into the recording head cap. The recording head cap was held by a multiaxis rotating arm (Instech: MLCA) at the top of the chamber to allow free movement of the rat during recording sessions.

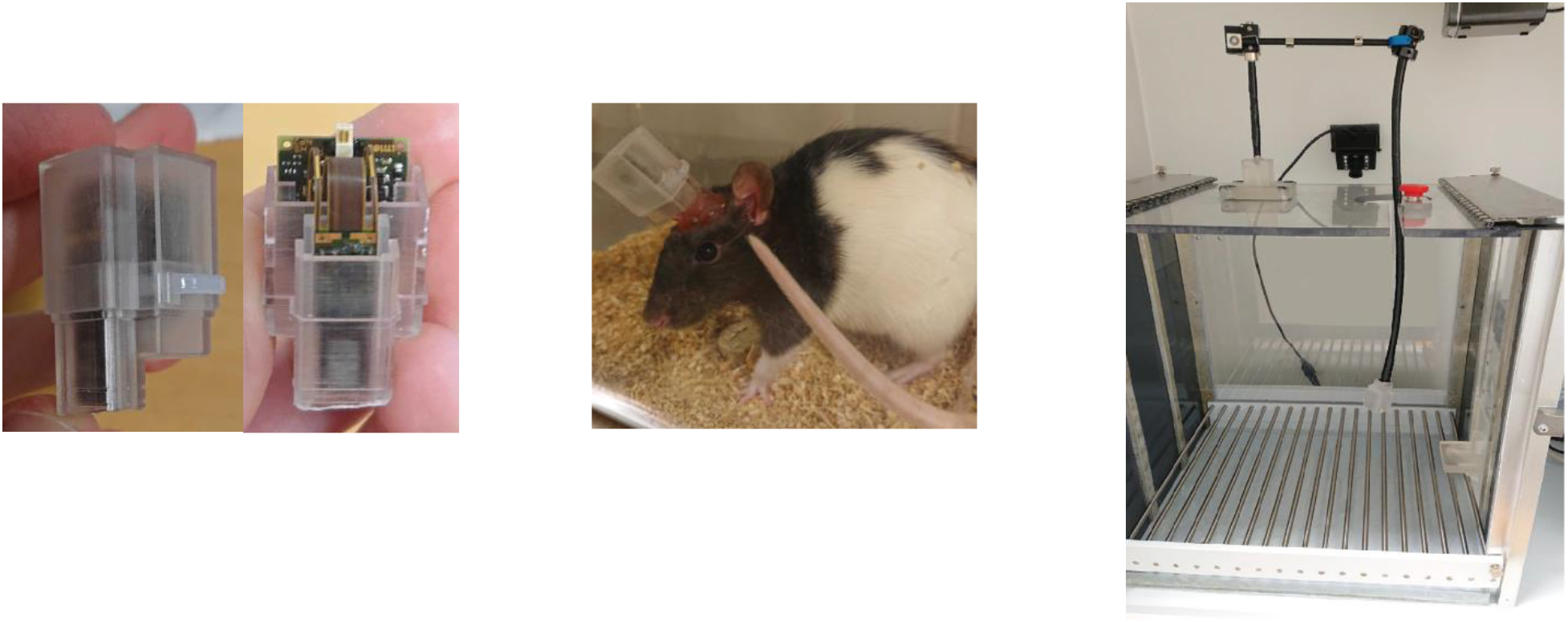

Prior to surgery vacuum grease was applied around the base of the front part of the head cap avoiding any grease touching the probe. The probe was then carefully painted with Dil (ThermoFisher, V22885). The skull was leveled, dried, and scored in a crosshatch pattern before four screws were fixed into the skull at locations leaving sufficient space for the head cap. A 1.4mm craniotomy was carried out and the dura removed. A clamp (southern labware: 30392204) was modified so that it could be used as a stereotaxic arm and hold the head cap. A piece of rubber tubing was put around the head cap to give the claw a better grip. The clamp holding the head cap was fixed to the stereotaxic arm at a 15⁰ angle, centered over the craniotomy, and slowly lowered until the probe came into contact with the cortex. The flat edge of a needle was carefully placed alongside the probe to prevent excessive bending and to help pierce the cortex. Once the probe visibly entered the cortex, it was slowly lowered until about half of the implant final depth was reached (∼3.5mm). Using a syringe and needle a small amount of the silicone compound (Dow: DOWSIL 3-4680) was injected into the craniotomy around the probe and given at least 5 minutes to set. The head cap was then lowered to its final position, either until the max depth was reached (7.5mm) or the head cap base came into contact with the skull.

Additional grease was applied around the base of the cap, sealing any gaps between the cap and the skull. The ground wire was wrapped around the two nearest skull screws in a figure of 8 pattern to ground the probe. The head cap was then cemented in place by filling the surrounding area with orthodontic resin (cc 22-05-98, Pearson Dental Supply), with care taken to ensure the cement reaches underneath the upper part of the head cap and around the entire base. A final amount of silicone gel was injected into the front part of the head cap to ensure the craniotomy was sealed. Once the cement and gel was set the clamp was released from around the head cap and the lid fixed in place with a cable tie.

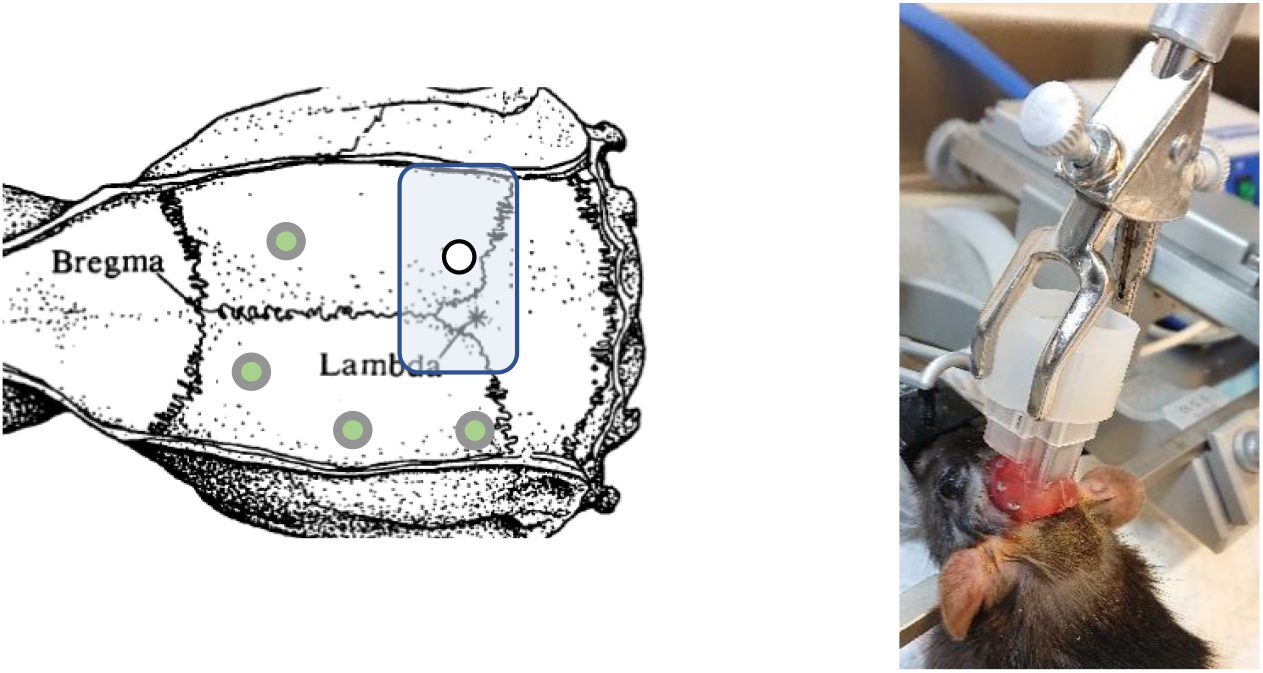

## Single unit data acquisition

An acquisition computer was connected to a PXIe-1071 chassis containing an acquisition module, a remote control module (PXIe-8381), and an I/O module (PXI-6363) with attached BNC connector block (2110). A med associates operant chamber was controlled using a separate computer running med-associates software. Behaviour data was collected using med-associates TTL boxes (SG-231) attached to the connector block. Single unit data was acquired using OpenEphys software with the Neuropix-PXI module (*1*). The NI-DAQmx module was used to record analogue signals of different lengths to timestamp events and behaviour of interest (*2*).

During each recording session the head cap lid was removed by cutting away the cable tie. The Neuropixels cable coming through the recording head cap was attached to the head stage and the recording lid secured with a new cable tie. The signal from the probe was checked in OpenEphys, and if successfully acquired without excessive noise or breaks in signal the recording channels for that session were selected in the Neuropix-PXI module. External or internal reference was selected to maximize the signal to noise ratio during recording. In the NI-DAQmx module all digital channels were unselected and the 3 analog channels for collecting behaviour data selected. The behaviour program was then started alongside recording in OpenEphys. Due to the large file size data were recorded directly to an external hard drive.

If signal was not acquired on first plugin in attempt the rat was re-plugged in. If signal was not able to be acquired after repeated plugin attempts signal was considered lost and recording session not carried out. If signal was still no longer able to be acquired the following day signal was considered lost and recording sessions ended for that subject. If signal was lost during the first ∼10 minutes of a session, the session was ended and re-started if signal was able to be re-acquired. If signal was lost further into the session, then the session was lost and recording ended for that subject for that day and resumed the following day. A camera inside the chamber allowed continued monitoring of the rat during the recording session. At the end of the recording session the rat was removed from the chamber and the lid but back onto the head cap with a new cable tie.

## Probe explant

After recording sessions were completed or terminated, the rat was put back under isoflurane anesthesia and returned to the stereotax in order to secure the head. The lid was removed, and the ground wire cut inside the head cap. Using a scalpel, the small dots of cement keeping the head stage in place were scraped off and the head stage removed. Next, the cement securing the probe mount was scraped off, and the mount very slowly pulled up/out of the head cap. Special care was taken not to pull too hard, otherwise the silicon in the bottom of the head cap may cause the probe to break. The retrieved probe was briefly soaked in DI water before being soaked overnight in a 1% tergazyme solution to remove residue silicon and biological material. Probes were then given a final rinse with DI water and 90% IPA before stored for future use.

## Identifying probe tracts

Immediately following probe retrieval rats were deeply anesthetized perfused intracardially and brains extracted. Extracted brains were frozen then sectioned for the full anterior posterior extent of the probe. Sections were mounted and processed with NeuroTrace, a cell body marker. Images of the full probe path were acquired with an Axioimager Z2 microscope. X/Y/Z coordinates were determined for the tip (most dorsal location) and top (most ventral location) of each probe implant. An implant vector was calculated based on these coordinates and each micrometer along the probe assigned x/y/z brain coordinates.

## Data processing and single unit sorting

All unit sorting and data analysis were carried on an analysis computer with a GeForce GTX 1080 GPU and 64GB RAM.

Data processing and analysis scripts will be uploaded and available at http://crcns.org/.

The aim of the unit sorting process was to ensure that each single unit accepted for analysis met the criteria of being an isolated single unit and was held for the entire 16 trials of the session in which it was recorded. Units were sorted using Kilosort2 (round 1 of recording) or Kilosort2.5 (round 2 of recording). Sorted sessions were loaded into Phy for manual curation process. Each unit identified by Kilosort was individually assessed in Phy using the ISI, correlogram, waveform, and feature view, and compared against other units for splits and merges. The temporal view was also examined to see if activity was recorded throughout the session or was dropped partway through. If a unit did not show clear separation from noise and/or other units, and/or did not show expected patterns of single unit activity (e.g. clear refractory period) it was not labelled as good and excluded from further analysis. On average 45.9 units per recording session were labelled as ‘good’ in phy and accepted for further analysis.

Using Matlab (with code from the Cortex-lab spikes repository (*3*)) information about firing and single unit location along the probe was extracted from the phy output for each session. Behaviour timestamps (danger/uncertainty/safety cues on, shock on, nose poke, pellet reward delivery) were extracted from the 3 analogue channels using a Matlab. A subsequent Matlab script created a single .mat file for each single unit that contained waveform time stamps, behaviour timestamps, and location on probe. Single units were then screened for stability across the recording session. Any unit that showed clear loss of activity during any of the trials was excluded from the analysis. Of the units that were still included at this stage, 52.6% met criteria to be included in the final analysis. During the screening stage it became apparent that there was a drift in the behaviour/events timestamps relative to the single unit firing data across each recording session. Using the timestamps recorded on the behaviour acquisition computer to compare to the timestamps extracted from the analogue data streams feeding into OpenEphys the drift was calculated and a correction applied to each single unit accepted for analysis (0.26s drift in the behaviour time stamps occurred during the 54-minute recording session).

Next the coordinates on the probe that each single unit was recorded from were extracted from the single unit Matlab files. For each subject the location that each single unit was recorded on the probe was mapped onto the coordinates calculated for that implant. These coordinates were then overlaid onto the rat atlas (*1*) in Matlab. Using these overlays and assigned coordinates each unit could then be manually assigned to a specific brain region. ‘Units’ obtained from the central aqueduct were removed. The final stage of unit processing was to run a cube script that compiled all the 1812 single unit Matlab files into a single array for analysis in Matlab.

## Z-score normalization

For each single unit and trial, firing rate (Hz) was calculated in 250 ms bins from 20 s prior to cue onset to 20 s following cue offset, for a total of 240 bins. Differential firing was calculated for each of the 240 bins by subtracting baseline firing rate (2 s prior to cue onset) specific to that trial type from each bin. Differential firing was *Z*-score normalized across all trials for each single unit, such that mean firing = 0, and standard deviation in firing = 1. *Z*-score normalization was applied to firing across the entirety of the recording epoch, as opposed to only the baseline period, in case single units showed little/no baseline activity. As a result, periods of marked firing increase during cue presentation would contribute to normalized mean firing rate (0). For this reason, *Z*-score normalized baseline activity can differ from zero.

## Heat plots

Heat plots were constructed from normalized firing rate using the imagesc function in Matlab. Mean normalized firing rate was calculated for each trial type for each single unit. For the cue heat map, this meant three trial types: danger, uncertainty, and safety. For the outcome heat map, this meant four trial types: danger, uncertainty shock, uncertainty omission, and safety. Perceptually uniform color maps were used to prevent visual distortion of the data (*2*).

## K-means clustering

For the cue period, brainstem single unit firing was organized into an 1812 single unit x 42-time interval matrix containing 14 x 1s intervals for each cue (1-14 danger, 15-28 uncertainty, 29-42 safety). These 14 intervals included the 2s pre-cue,10s cue, and 2s following cue offset but before shock delivery. Clustering was carried out using the Matlab kmeans function. K-means clustering results were number identifier for each single unit and the mean of the squared Euclidean distance from cluster centroid. K-means clustering was performed 25 times, incrementing the number of clusters from 1-25. For each iteration we compared the number of clusters, the mean of the squared Euclidian distance and the number of single units composing the clusters. These three factors were examined for over and under clustering. Over clustering was apparent when clusters contained only a single or low number of single units, and when adding additional clusters marginally reduced the mean squared Euclidean distance. Under clustering was apparent when clusters had large mean squared Euclidean distances and dissimilar single unit types were lumped together. The ideal # of clusters for each firing period found the midpoint of these extremes. K-means clustering was performed identically for the outcome period, only now firing data came from the post shock period. An 1812 single unit x 80 time-interval matrix was constructed from the 20, 500ms intervals for each trial type 10s from shock offset (1-20 danger, 21-40 uncertainty shock, 41-60 uncertainty shock omission, 61-80 safety). K-means clustering was performed 20 times, incrementing the number of clusters from 1-20.

## Principal components analysis

Principal component analysis (PCA) was performed to reveal low-dimensional brainstem firing features. Brainstem single-unit firing was organized into an 1812 single unit x 42 column matrix (14, 1s intervals for each trial type) for the cue period; and an 1812 x 80 column matrix (20, 500ms intervals for each trial) matrix for the outcome period. PCA was performed using the pca function in Matlab. PCA outputs were the PC weights (cue x time) for each principal component as well as the percent firing variance explained for each principal component (Fig 2C).

Cluster identity from k-means clustering, principal components analysis combined with iterative shuffling to determine each clusters contribution to principal component 1. First, the percent firing variance explained by PC1 was determined for all brainstem single units as described above. This result served as the ground truth reference for subsequent results. Next, a single cluster was selected (i.e., cluster 21). Firing information for the single units of only that cluster were shuffled. Firing information for all other clusters (i.e., clusters 1-20) was left intact. Principal components analysis was performed on this modified data set (clusters 1-20 intact, cluster 21 shuffled) and the percent firing variance explained for PC1 determined. PC1 contribution for that iteration was determined by subtracting the modified data set (clusters 1-20 intact, cluster 21 shuffled) from the ground truth (all clusters intact). Shuffling → PCA was performed 1,000 times for each cluster, and PC1 contribution determined for each iteration, then averaged. The complete iterative, shuffle/PCA was performed for each individual cluster, always shuffling the cluster of interest but leaving intact all other clusters. Clusters more greatly contributing to PC1 would have larger difference values, indicating that removing their firing information more greatly reduced the percent of firing variance explained by PC1.

## Danger firing latency

Mean normalized firing rate during 10-s danger cue presentation was summarized in 10,000, 1ms bins for each single unit. Danger firing latency was determined for each single unit by identifying the bin of greatest firing inflection (among the 10,000 total during danger presentation) using the findchangepts Matlab function (Fig 2D). Mean danger firing latencies were calculated for each cluster by averaging the latencies of the composing single units. Between-cluster differences in latency means were determined using Bonferroni-corrected (p = 0.05/# of tests), independent samples t-tests. Between-cluster differences in latency variation were determined using Bonferroni-corrected, Levene’s test for equality of variances.

## Correlated cluster firing

Mean cue firing for each cluster was organized into an 1812 single unit x 36 column matrix (12, 1s intervals for each cue, 2s pre-cue through 10s cue). Mean cue firing for each cluster was correlated with mean cue firing for every other cluster. The result was a 21 x 21 matrix of the Pearson’s correlation coefficient (R^2^) for each cluster comparison (Fig 2E; 4D). Correlation coefficients on the diagonal were always 1, as they were a cluster’s correlation with itself. Correlated cluster firing was also performed for the outcome period, for which cluster outcome firing was organized into an 1812 single unit x 80 column matrix (20, 500ms intervals for each trial type, 10s following shock offset).

## Hub identification with single unit x cluster correlations

For each single unit from each of the eight cue subnetwork clusters (i.e., cluster 1), single-unit firing during the danger cue (14 x 1s intervals, starting 2s pre-cue) was correlated against mean danger firing of each remaining seven clusters (i.e., clusters 2-8). The result was seven separate correlation coefficients for each single unit (R^2^), one for each cluster comparison. A mean correlation coefficient was calculated for each single unit, providing a measure of how well each unit correlated with the firing of the other subnetwork clusters (Fig 2F). Independent samples t-tests were used to determine differences between-cluster differences in mean R^2^. Levene’s test for equality of variance was used to determine differences between-cluster differences in standard deviation of R^2^. Bonferroni correction was applied to both comparisons, producing a significance threshold of 0.007 (0.05/7). A cluster showing the highest mean and least variable R^2^ was considered a hub. Similar analyses were performed for the phasic outcome subnetwork (clusters 1-5) and the tonic outcome subnetwork (clusters 7-10); only now, firing was taken from all trials types (20, 500ms intervals for each trial type, 10s following shock offset).

| Trial # | Trial type | Z firing | Constant | Threat probability | Behaviour |
| --- | --- | --- | --- | --- | --- |
| 1 | danger | 1.49 | 1 | 1 | 0.98 |
| 2 | danger | 1.89 | 1 | 1 | 0.85 |
| 3 | danger | 2.42 | 1 | 1 | 0.70 |
| 4 | danger | 2.04 | 1 | 1 | 0.85 |
| 5 | uncertainty | 1.16 | 1 | 0.25 | 0.45 |
| 6 | uncertainty | 1.20 | 1 | 0.25 | 0.79 |
| 7 | uncertainty | 1.32 | 1 | 0.25 | 0.79 |
| 8 | uncertainty | 1.63 | 1 | 0.25 | 0.77 |
| 9 | uncertainty | 1.02 | 1 | 0.25 | 0.64 |
| 10 | uncertainty | 1.18 | 1 | 0.25 | 0.71 |
| 11 | uncertainty | 1.19 | 1 | 0.25 | 0.55 |
| 12 | uncertainty | 0.96 | 1 | 0.25 | 0.48 |
| 13 | safety | 0.77 | 1 | 0 | 0.46 |
| 14 | safety | 0.93 | 1 | 0 | 0.41 |
| 15 | safety | 0.57 | 1 | 0 | 0.23 |
| 16 | safety | 0.52 | 1 | 0 | 0.30 |
| Beta coefficient |  |  |  | 0.97 | 0.49 |

## Linear regression

Linear regression (example shown above) was used to determine whether cue cluster firing was better explained by threat probability or behaviour. Separately for each of the 21 cue clusters, firing was organized by trial type (16 trials, 3 types) over 14, 1s intervals (2s pre-cue through 2s post-cue). The threat probability regressor was the foot shock probability associated with each cue (1 or 0.25 or 0). The behaviour regressor was the trial-specific, mean suppression ratio for the cluster. The Matlab regress function was used simultaneously compare which regressor better explained cluster firing for each interval. Regression output was a beta coefficient for each regressor and interval (fig 3A). A beta coefficient quantified the direction (>0 = positive) and strength (>0 = stronger) of the relationship between each regressor and cluster firing.

Regression was also performed for the 11 outcome clusters. Firing was organized by trial type (16 trials, 4 types) over 20, 500ms intervals (10s following shock offset). The regressors were shock: 1 (danger), 1 (uncertainty shock), 0 (uncertainty omission), and 0 (safety); and prediction error: 0 (danger), 0.75 (uncertainty shock), −0.25 (uncertainty omission), and 0 (safety). Regression output was a beta coefficient for each regressor and interval. A beta coefficient quantified the direction (>0 = positive) and strength (>0 = stronger) of the relationship between each regressor and cluster firing.

## ‘Lesion’ x principal components analysis x linear regression

Principal component analysis (PCA) was performed to reveal low-dimensional brainstem signaling features. Cluster beta coefficients for threat probability and behaviour were organized into a 21-cluster x 28-interval matrix (intervals 1-14 probability, 15-28 behaviour) for the cue period. PCA was performed using the pca function in Matlab. PCA outputs were the weights (regressor x time) for each principal component (we focus on PC1 and PC2) as well as the percent signaling variance explained by PC1 and PC2 (Fig 2C). PCA was first performed with all clusters and networks ‘intact’. PCA was similar for the outcome period, only now cluster beta coefficients for signed prediction error and shock were organized into a 21-cluster x 40-interval matrix (intervals 1-20 shock, 21-40 prediction error) for the post-shock period.

The ‘lesion’ analysis was similar to that used for determining the contribution of each cluster to low dimensional firing features. Only now, the lesion approach was used to determine differential contributions of the cue subnetwork and supranetwork to low dimensional cue signaling; as well as differential contributions of the phasic and tonic outcome subnetworks to low dimensional outcome signaling.

First, the PC1 and PC2 weight patterns (regressor x time) were determined for all 21 clusters as described above. Next, subnetwork clusters 1-8 were selected. Beta coefficients for only these cluster were shuffled. Principal components analysis was performed on this modified data set (clusters 1-8 shuffled, clusters 9-21 intact) and the PC1 and PC2 weight patterned determined. Shuffling → PCA was performed 1,000 times, and PC1/PC2 weights determined for each iteration, then averaged for all 1,000 iterations. All signaling by the cue subnetwork would be lost, while signaling by the supranetwork would be intact. The inverse analysis was performed to determine signaling by the supranetwork: subnetwork clusters were intact but supranetwork clusters were lesioned. Finally, the outcome ‘lesion’ analysis contrasted contribution of the phasic and tonic outcome networks by shuffling one network, while leaving the other intact. Comparing PC1 and PC2 weights for the fully intact regression x PCA versus phasic shuffled and tonic shuffled allowed to reveal the specific contribution of each network to outcome signaling.

## Mapping cluster and network membership to brain region

For each cue and outcome cluster, the percent of member single units originating from each brain region was determined and plotted using the Matlab bubchart function (Fig. 2G; 4G).

To map network membership to brain region, the percent of single units from each region contributing to the subnetwork vs. supranetwork and phasic vs. tonic outcome network were calculated and visualized with donut charts using the donut function in Matlab. To determine the differential contribution of each region to functional networks, first the population proportions of subnetwork vs. supranetwork and phasic vs. tonic outcome network were determined. Then for each brain region, the proportion contributing to each network was subtracted from the overall proportion. Values around zero would indicate a distribution of subnetwork vs. supranetwork and phasic vs. tonic outcome network clusters that reflected the entire brainstem. Deviations from zero would indicate preferential network contributions.

## Determining temporal firing pattern following shock

Temporal firing pattern following foot shock was calculated by averaging normalized z firing following shock on danger and uncertainty trials. Differential firing was then calculating mean normalized firing 1s following shock offset – mean normalized firing 5s following shock offset. The absolute value was taken to fairly compare units increasing vs. decreasing firing following foot shock. Phasic neurons will have high values, while tonic neurons will have low values.

## Relationship between cue network and outcome network membership

Chi square testing was used to determine the relationship between cue subnetwork membership and outcome network membership. 505/1812 (27.9%) single units were in the cue subnetwork. Therefore, chi-square test determined if the proportion of phasic outcome single units in the cue subnetwork differed from 27.9%. The same was asked for the tonic outcome single units. Results were a chi-squared statistic and associated p value.

## Notes

### Competing Interest Statement

The authors have declared no competing interest.

### Summary of Updates

We reviewed our firing analyses for our manuscript and uncovered an error in trial averaging. Our discrimination sessions consist of 16 trials (4 danger, 2 uncertainty shock, 6 uncertainty omission, and 4 safety). Trial type order is randomized so that every session is unique. To analyze firing and behavior across all sessions, we must first standardize the trial order. For each of the 16 session trials we determine the type (of 4 possible) and the number of occurrences for that type. We then arrange the trial types in a standardized order for analysis. Danger trials are first, uncertainty shock second, uncertainty omission third, and safety last. Trial number is maintained within each trial type. Danger trial 1 is first, trial 2 second, etc. We perform this standardization for every session, which results in a 3D matrix: trial types are rows (X), time bins are columns (Y), and sessions are stacked layers (Z). Many analyses require us to average all trials of a single type. This is where the error occurred. To find mean danger firing, we average rows 1:4. For uncertainty shock, rows 5:6; uncertainty omission, rows 7:12; and for safety, rows 13:16. However, when we checked our code a 1 was omitted from safety averaging. In error, we averaged rows 3:16 instead of 13:16. What we thought was mean safety firing was actually mean firing for ALL cues, except the first two danger cues. The error has been fixed. The revised manuscript uses the accurate safety firing data.

## References

1. I. Levy, D. Schiller, Neural Computations of Threat. Trends Cogn Sci. 25, 151–171 (2021).

2. M. S. Fanselow, Neural Organization of the Defensive Behavior System Responsible for Fear. Psychon B Rev. 1, 429–438 (1994).

3. T. Ozawa, E. A. Ycu, A. Kumar, L. F. Yeh, T. Ahmed, J. Koivumaa, J. P. Johansen, A feedback neural circuit for calibrating aversive memory strength. Nat Neurosci. 20, 90–97 (2017).

4. P. Tovote, M. S. Esposito, P. Botta, F. Chaudun, J. P. Fadok, M. Markovic, S. B. Wolff, C. Ramakrishnan, L. Fenno, K. Deisseroth, C. Herry, S. Arber, A. Luthi, Midbrain circuits for defensive behaviour. Nature. 534, 206–12 (2016).

5. K. M. Wright, M. A. McDannald, Ventrolateral periaqueductal gray neurons prioritize threat probability over fear output. Elife. 8 (2019), doi:10.7554/eLife.45013.

6. K. M. Wright, T. C. Jhou, D. Pimpinelli, M. A. McDannald, Cue-inhibited ventrolateral periaqueductal gray neurons signal fear output and threat probability in male rats. Elife. 8 (2019), doi:10.7554/eLife.50054.

7. R. A. Walker, K. M. Wright, T. C. Jhou, M. A. McDannald, The ventrolateral periaqueductal gray updates fear via positive prediction error. Eur J Neurosci (2019), doi:10.1111/ejn.14536.

8. M. Roy, D. Shohamy, N. Daw, M. Jepma, G. E. Wimmer, T. D. Wager, Representation of aversive prediction errors in the human periaqueductal gray. Nature neuroscience. 17, 1607–12 (2014).

9. R. A. Rescorla, A. R. Wagner, in Classical Conditioning II: Current Research and Theory, B. AH, P. WF, Eds. (Appleton Century Crofts, New York, 1972), pp. 64–99.

10. J. J. Jun, N. A. Steinmetz, J. H. Siegle, D. J. Denman, M. Bauza, B. Barbarits, A. K. Lee, C. A. Anastassiou, A. Andrei, C. Aydin, M. Barbic, T. J. Blanche, V. Bonin, J. Couto, B. Dutta, S. L. Gratiy, D. A. Gutnisky, M. Hausser, B. Karsh, P. Ledochowitsch, C. M. Lopez, C. Mitelut, S. Musa, M. Okun, M. Pachitariu, J. Putzeys, P. D. Rich, C. Rossant, W. L. Sun, K. Svoboda, M. Carandini, K. D. Harris, C. Koch, J. O’Keefe, T. D. Harris, Fully integrated silicon probes for high-density recording of neural activity. Nature. 551, 232–236 (2017).

11. M. Moaddab, M. A. McDannald, Retrorubral field is a hub for diverse threat and aversive outcome signals. Current Biology. 31, 2099–2110.e5 (2021).

12. B. A. Berg, G. Schoenbaum, M. A. McDannald, The dorsal raphe nucleus is integral to negative prediction errors in Pavlovian fear. European Journal of Neuroscience. 40, 3096– 3101 (2014).

13. G. Paxinos, C. Watson, The rat brain in stereotaxic coordinates (Academic Press/Elsevier, Amsterdam ; Boston ;, ed. 6th, 2007; Publisher description http://www.loc.gov/catdir/enhancements/fy0745/2006937142-d.html).

14. J. P. Johansen, J. W. Tarpley, J. E. LeDoux, H. T. Blair, Neural substrates for expectation-modulated fear learning in the amygdala and periaqueductal gray. Nature neuroscience. 13, 979–86 (2010).

15. A. A. Grace, Phasic versus tonic dopamine release and the modulation of dopamine system responsivity: A hypothesis for the etiology of schizophrenia. Neuroscience. 41, 1–24 (1991).

16. K. M. Rothenhoefer, T. Hong, A. Alikaya, W. R. Stauffer, Rare rewards amplify dopamine responses. Nat Neurosci. 24, 465–469 (2021).

17. M. Alexandra Kredlow, R. J. Fenster, E. S. Laurent, K. J. Ressler, E. A. Phelps, Prefrontal cortex, amygdala, and threat processing: implications for PTSD. Neuropsychopharmacol., 1–13 (2021).

18. S. Duvarci, D. Pare, Amygdala microcircuits controlling learned fear. Neuron. 82, 966–80 (2014).

19. K. Kveraga, J. Boshyan, R. B. Adams, J. Mote, N. Betz, N. Ward, N. Hadjikhani, M. Bar, L. F. Barrett, If it bleeds, it leads: separating threat from mere negativity. Soc Cogn Affect Neur. 10, 28–35 (2015).

20. B. J. Liddell, K. J. Brown, A. H. Kemp, M. J. Barton, P. Das, A. Peduto, E. Gordon, L. M. Williams, A direct brainstem-amygdala-cortical “alarm” system for subliminal signals of fear. Neuroimage. 24, 235–243 (2005).

21. J. M. P. Baas, J. Milstein, M. Donlevy, C. Grillon, Brainstem Correlates of Defensive States in Humans. Biological Psychiatry. 59, 588–593 (2006).

22. P. A. Kragel, M. Čeko, J. Theriault, D. Chen, A. B. Satpute, L. W. Wald, M. A. Lindquist, L. Feldman Barrett, T. D. Wager, A human colliculus-pulvinar-amygdala pathway encodes negative emotion. Neuron. 109, 2404–2412.e5 (2021).

23. Y. C. Wang, M. Bianciardi, L. Chanes, A. B. Satpute, Ultra High Field fMRI of Human Superior Colliculi Activity during Affective Visual Processing. Sci Rep. 10, 1331 (2020).

24. Z. Zhao, M. Davis, Fear-Potentiated Startle in Rats Is Mediated by Neurons in the Deep Layers of the Superior Colliculus/Deep Mesencephalic Nucleus of the Rostral Midbrain through the Glutamate Non-NMDA Receptors. J. Neurosci. 24, 10326–10334 (2004).

25. O. K. Faull, M. Jenkinson, M. Ezra, Kt. Pattinson, Conditioned respiratory threat in the subdivisions of the human periaqueductal gray. Elife. 5 (2016), doi:10.7554/eLife.12047.

26. A. B. Satpute, T. D. Wager, J. Cohen-Adad, M. Bianciardi, J. K. Choi, J. T. Buhle, L. L. Wald, L. F. Barrett, Identification of discrete functional subregions of the human periaqueductal gray. P Natl Acad Sci USA. 110, 17101–17106 (2013).

27. G. P. McNally, S. Cole, Opioid receptors in the midbrain periaqueductal gray regulate prediction errors during pavlovian fear conditioning. Behavioral neuroscience. 120, 313–23 (2006).

28. K. E. Sos, M. I. Mayer, C. Cserép, F. S. Takács, A. Szőnyi, T. F. Freund, G. Nyiri, Cellular architecture and transmitter phenotypes of neurons of the mouse median raphe region. Brain Struct Funct. 222, 287–299 (2017).

29. A. Ploghaus, I. Tracey, S. Clare, J. S. Gati, J. N. P. Rawlins, P. M. Matthews, Learning about pain: The neural substrate of the prediction error for aversive events. P Natl Acad Sci USA. 97, 9281–9286 (2000).

30. V. I. Spoormaker, K. C. Andrade, M. S. Schroter, A. Sturm, R. Goya-Maldonado, P. G. Samann, M. Czisch, The neural correlates of negative prediction error signaling in human fear conditioning. Neuroimage. 54, 2250–2256 (2011).

31. W. Schultz, P. Dayan, P. R. Montague, A neural substrate of prediction and reward. Science. 275, 1593–9 (1997).

32. M. R. Roesch, D. J. Calu, G. Schoenbaum, Dopamine neurons encode the better option in rats deciding between differently delayed or sized rewards. Nature Neuroscience. 10, 1615– 24 (2007).

33. N. Assareh, E. E. Bagley, P. Carrive, G. P. McNally, Brief optogenetic inhibition of rat lateral or ventrolateral periaqueductal gray augments the acquisition of Pavlovian fear conditioning. Behav Neurosci. 131, 454–459 (2017).

34. L.-F. Yeh, T. Ozawa, J. P. Johansen, Functional organization of the midbrain periaqueductal gray for regulating aversive memory formation. Mol Brain. 14, 136 (2021).

35. M. H. Ray, E. Hanlon, M. A. McDannald, Lateral orbitofrontal cortex partitions mechanisms for fear regulation and alcohol consumption. PLoS One. 13, e0198043 (2018).

36. C. H. Beck, H. C. Fibiger, Conditioned fear-induced changes in behavior and in the expression of the immediate early gene c-fos: with and without diazepam pretreatment. Journal of Neuroscience. 15, 709–20 (1995).

37. G. Vetere, J. W. Kenney, L. M. Tran, F. Xia, P. E. Steadman, J. Parkinson, S. A. Josselyn, P. W. Frankland, Chemogenetic Interrogation of a Brain-wide Fear Memory Network in Mice. Neuron. 94, 363–374 e4 (2017).

38. D. V. P. S. Murty, S. Song, K. Morrow, J. Kim, K. Hu, L. Pessoa, Distributed and Multifaceted Effects of Threat and Safety. Journal of Cognitive Neuroscience. 34, 495–516 (2022).

39. M. Pachitariu, N. Steinmetz, S. Kadir, M. Carandini, K. Harris, Fast and accurate spike sorting of high-channel count probes with KiloSort. Adv Neur In. 29 (2016) (available at ://WOS:000458973702073).

40. F. Crameri, Scientific colour maps (Version 4.0.0). (2018),, doi:10.5281/zenodo.2649252.

## References

1. Siegle, J., Kulik, P., & Doshi, A. Neuropixels PXI plugin. https://github.com/open-ephys-plugins/neuropixels-pxi (2019).

2. Kulik, P. NI-DAQ plugin. https://github.com/open-ephys-plugins/nidaq-plugin (2019).

3. Steinmetz, N. spikes. https://github.com/cortex-lab/spikes/ (2019).

4. F. Crameri, Scientific colour maps (Version 4.0.0). (2018), doi:10.5281/zenodo.2649252.

